# In Vitro Detection of Breast Cancer Cell Types Using Machine Learning-Assisted Spectral Fingerprinting of SWCNTs

**DOI:** 10.64898/2026.06.30.735651

**Authors:** Maryam Rahmani, Katherine Van Gorden, Shelly R. Peyton, Daniel Roxbury

## Abstract

The early detection of breast cancer currently relies on expensive mammography, followed by pathology that uses biopsied, fixed, and immunohistochemically stained tissues. A live-cell detection approach could be highly beneficial as a supportive diagnostic and research tool to better understand and resolve the dynamic nature of breast cancer cells and their response to treatment in real time. Here, we present a single-walled carbon nanotube (SWCNT) near-infrared fluorescence spectral fingerprinting approach combined with machine learning to precisely detect the heterogeneity of breast cancer cells in live culture. We introduced DNA-functionalized SWCNTs to MCF-10A (a non-tumorigenic healthy control) and cancer cell lines spanning known extrinsic disease subtypes: MCF-7 (luminal A), HCC1954 (HER2+), MDA-MB-231, and MDA-MB-468 (both triple-negative). The NIR fluorescence spectra of DNA-SWCNTs across 600 individual cells within each type showed significant differences in emission peak intensities, center wavelengths, and peak intensity ratios, attributable to variations in cellular uptake and biomolecular interactions. These spectral changes likely arise from complex SWCNT–cellular interaction “fingerprint” that includes redox-mediated modulation of the local nanotube environment, rather than from a single biomarker response. The extracted spectral features were used to train an ensemble machine learning model. The model achieved 98% classification accuracy for breast cancer detection and 95% classification accuracy for breast cancer cell subtyping. Moreover, Raman microscopy further showed that MDA-MB-468 cells exhibited the highest SWCNT uptake, whereas MCF-10A cells showed greater SWCNT aggregation, consistent with their lower broadband NIR fluorescence intensity. These results demonstrate that SWCNT NIR fluorescence fingerprints can capture cell line-specific optical signatures. This platform provides a foundation for nanomaterial-enabled biosensing strategies aimed at real-time monitoring of cancer-associated cellular states.

## Introduction

Breast cancer is the most commonly diagnosed cancer in women worldwide and remains a leading cause of cancer related death. In 2022, approximately 2.3 million new cases and 670,000 deaths were reported globally, and those numbers are projected to keep rising, with annual cases expected to exceed 3.2 million new cases and 1.1 million breast cancer-related deaths by 2050.^1,2^ Deaths relative to new cases range from around 18% in highly developed countries to over 50% in less developed countries, with a portion of this decrease related to the availability of early detection techniques. This discrepancy proves that access to early detection diagnostic techniques and proper treatments are vital to a patient’s survival.^2^

Breast tumors can be classified into intrinsic and extrinsic subtypes, and extrinsic subtyping performed by pathologists is used by oncologists to plan treatment strategies^3–6^. These subtypes are classified based on expression of specific receptors, and include luminal A (estrogen and progesterone receptor-positive, human epithelial growth factor negative), luminal B (estrogen or progesterone receptor-positive), HER2+ (human epithelial growth factor receptor 2)-positive, and triple negative (negative for all three hormone receptors).^7,8^ Luminal A subtype is the most common, accounting for approximately 68% of breast cancer cases in the United States.^8^ It generally has the best prognosis, driven by hormone receptor positivity and responsiveness to endocrine therapies.^9,10^ The HER2-Positive subtype accounts for 5% of cases in the US, it overexpresses the HER2 receptor, and is more aggressive than the luminal subtypes if left untreated. HER2-targeting agents such as Herceptin, trastuzumab, and newer antibody drug conjugates such as trastuzumab emtansine and trastuzumab deruxtecan are effective against HER2+ breast tumors.^11–13^ Triple negative cases account for roughly 10% of breast cancers^8^, have no hormone-targeted therapy, and are associated with early relapse and poor survival after metastasis.^14,15^

Detecting breast cancer early significantly improves outcomes. The 5-year relative survival rate at a localized stage is over 99%, and drops to 32% at a distant stage^13^, but existing screening and subtyping tools have real limitations. Mammography, the universal current method for screening in developed nations, has false negative rates of 10-12% and has a decreased performance in women with dense breast tissue.^16,17^ When a tumor is detected, accurately classifying its subtype to guide treatment requires immunohistochemistry (IHC) and analysis by a pathologist in fixed biopsy tissue. This method is invasive and static, meaning they only capture receptor expression at a single point in time and requires tissue fixation^18^, which makes them unfit for monitoring dynamic changes in cellular state. This creates a clear need for new platforms that can identify breast cancer cell types reliably, ideally in live cells.

Single-walled carbon nanotubes (SWCNTs), an effectively one-dimensional carbon-based nanomaterial, have demonstrated applications as intracellular optical biosensors.^19–21^ Semiconducting SWCNTs emit fluorescence in the near-infrared (NIR) range of 900-1600 nm, a spectral window where biological tissue exhibits low autofluorescence, absorption, and scattering, making it desirable for biological imaging.^22,23^ Unlike organic fluorophores, SWCNTs do not photobleach, rather they can be imaged continuously over long periods of time without signal degradation.^23^ SWCNTs are produced as a family of species (chiralities), designated by their distinct (*n*,*m*) chiral indices. Each SWCNT chirality emits narrow-bandwidth fluorescence at bandgap-dependent wavelengths in the NIR, enabling multiplexed output from a single sample.^24–26^ SWCNT emission is sensitive to the local nanoscale environment, including changes in pH, dielectric environment, aggregation state, and biomolecular adsorption, which all produce measurable shifts in NIR fluorescence peak position, intensity, and bandwidth.^27–31^ Specific SWCNT-based sensors have been designed to detect target proteins^32^, lipids^33^, and neurotransmitters^34,35^ such as dopamine. Raman microscopy complements this by providing quantitative information about SWCNT concentration and structural integrity. The G-band (1585 cm^-1^) scales linearly with nanotube concentration, while the ratio of the disorder induced D-band (1350 cm^-1^) to the G-band shows structural defects that can be introduced during cellular processing.^36,37^

To enter live cells, SWCNTs must be properly dispersed and functionalized. Noncovalent functionalizations are preferred in order to preserve the desirable optical properties of the SWCNTs. Short strands of single stranded DNA (ssDNA), i.e. an amphiphilic biomolecule, is routinely used to disperse SWCNTs and impart biocompatibility to the resulting DNA-SWCNT hybrids.^38,39^ Upon introduction to live cells, DNA-SWCNTs are internalized through energy dependent endocytosis and progress through the endolysosomal pathway, eventually settling in lysosomes.^33,40^ As the DNA-SWCNTs translocate between intracellular vesicles, the local environment varies significantly across cell types in terms of pH, ion composition, protein content, and enzyme activity, and those differences are reported in the optical response of the internalized DNA-SWCNTs.^31^ We have previously shown that these variations can be used to benchmark the process of endocytosis as well as phenotype live macrophages in a machine learning-assisted technique that we termed SWCNT spectral fingerprinting.^41^ In this approach, high-dimensional NIR spectral datasets are utilized, necessitating machine learning algorithms to extract meaningful classification information.

Broadly speaking, approaches like convolutional neural networks, random forests, and ensemble methods have become standard tools in biomedical imaging.^42,43^ Specifically, optimizable ensemble models reduce prediction variance and bias while improving model stability, resulting in more robust classification of complex, high-dimensional datasets.^44^ Since biological data are often high-dimensional, noisy, nonlinear, correlated, and limited in sample size,^25^ ensemble methods have demonstrated strong empirical performance in this area (e.g., gene expression classification,^45^ single-cell Raman discrimination,^46^ and cancer detection^47,48^). AdaBoost (Adaptive Boosting), introduced by Freund and Schapire in 1997, is an ensemble learning algorithm that combines multiple weak classifiers sequentially, with each iteration focusing on the samples misclassified by the previous one.^49^ This results in a strong classifier that handles high-dimensional data well, even with relatively small training set sizes. Ensemble methods like AdaBoost have been used successfully for a range of medical classification tasks, including breast MRI analysis and Alzheimer’s diagnosis.^50,51^

Here, utilizing machine learning-assisted SWCNT spectral fingerprinting on DNA-SWCNTs internalized within live cells, we distinguish four different epithelial breast cancer cell lines versus a healthy control (MCF10A). The cancer cell lines span multiple subtypes: MCF-7 (luminal A), HCC1954 (HER2+), MDA-MB-231, and MDA-MB-468 (both triple negative) and MCF10A (healthy control) cell line. An ensemble algorithm machine learning model was trained on the fluorescence spectral responses from the 5 cell lines in order to detect breast cancer with a high accuracy of 98% and differentiate the cell lines from each other with a high accuracy of 95%. Raman microscopy was used to compare SWCNT uptake across cell lines. DNA-SWCNTs were internalized by all cell lines without inducing significant cytotoxicity, with the highest uptake observed in MDA-MB-468 cells. In contrast, SWCNTs exhibited the greatest degree of intracellular aggregation in MCF10A cells. These findings support the development of SWCNT based nanomaterial platforms^25^ for precision oncology and provide a foundation for future real time monitoring of breast cancer cells.

## Material and method

### SWCNT Preparation

Mono-dispersed ssDNA-wrapped SWCNTs (DNA-SWCNTs) were prepared with a mass ratio of 1:2 of SWCNTs: DNA by adding 1 mg of raw HiPCO SWCNT powder to 2 mg of (GT)_6_ oligonucleotide (Integrated DNA Technologies) and 1 ml of 0.1 M NaCl (Sigma-Aldrich). DNA functionalized SWCNTs were prepared using 1/8′′ tapered microtip and ultrasonicated for 30 min at 40% amplitude in an ice bath (Sonics Vibracell VCX-130; Sonics and Materials) and ultracentrifuged (Beckman Optima MAX-XP) for 30 min at 60,000 rpm and 10 °C. The top ∼80% of the supernatant was then collected to separate the aggregations. The concentration of DNA-SWCNTs was determined by acquiring a UV-Vis-NIR absorbance scan (Jasco, Tokyo, Japan) and utilizing an extinction coefficient of A_910_ = 0.02554 L mg^-1^cm^-1^.^27^

### Cell culturing

Human breast epithelial cell lines, including the non-tumorigenic control MCF-10A, and breast cancer cell lines MCF-7 (luminal), HCC1954 (HER2+), MDA-MB-231 (triple-negative, claudin-low), and MDA-MB-468 (triple-negative, basal-like) (ATCC, Manassas, VA, USA), were cultured under standard conditions at 37 °C in a humidified atmosphere containing 5% CO_2_. MCF-7, MDA-MB-231, and MDA-MB-468 cells were maintained in high-glucose Dulbecco’s Modified Eagle Medium (DMEM), while HCC1954 cells were cultured in Roswell Park Memorial Institute 1640 (RPMI 1640) medium. Both DMEM and RPMI media were supplemented with 10% fetal bovine serum (FBS), 2.5% HEPES, 1% L-glutamine, and 1% penicillin/streptomycin. MCF-10A cells were cultured in Mammary Epithelial Basal Medium (MEBM; Lonza) supplemented with 10% donor horse serum (DHS) and the SingleQuots TM supplement kit (Lonza). MCF-7, HCC1954, MDA-MB-231, and MDA-MB-468 cells were passaged regularly using trypsinization. Briefly, the culture medium was removed, the cells were rinsed with PBS (2x, 2 mL per wash), then incubated with 3 mL of trypsin at 37 °C for 7 min to facilitate detachment and reseeded into new 75 cm^2^ cell culture flasks. MCF-10A cells were passaged following the ATCC-recommended protocol using 0.05% trypsin–0.02% EDTA. Briefly, the culture medium was removed, and the cell monolayer was rinsed with DPBS (2x, 2 mL per wash). Then, 3 ml of trypsin–EDTA was added to the 75 cm^2^ cell culture flask, and the flask was incubated at 37 °C for 15 min to detach the cells. Trypsin activity was neutralized with 3 ml of 0.1% soybean trypsin inhibitor in DPBS, and the cell suspension was subsequently centrifuged (400×g, 10 min). The resulting cell pellet was resuspended in complete growth medium, reseeded into new culture vessels, and used for experiments.

### Near-Infrared Fluorescence Imaging

Hyperspectral near-infrared (NIR) fluorescence imaging was performed to obtain spectral fingerprints from individual cells following SWCNT internalization. Cells were imaged using a hyperspectral NIR fluorescence Olympus IX-73 inverted microscope with a 730 nm excitation laser source, LCPlan N, 20×/0.45 IR objective by Olympus, U.S.A. Hyperspectral image stacks were produced by passing the emitted fluorescence through a volume Bragg grating and detecting it with a 2D InGaAs array detector (Photon Etc., Montreal, Canada). In vitro imaging was performed under standard cell culture conditions at 37 °C and 5% CO_2_ in an Okolab stage-top incubator. Fluorescence images and hyperspectral cubes were collected using integration times of 0.2s and 1.5s and then background-subtracted using custom MATLAB scripts.

For all 20× in vitro NIR fluorescence imaging experiments, cells were plated at an initial concentration of 3×10^5^ cells/cm^2^ in 35mm non-glass-bottom microwell dishes (CellTreat) and cultured overnight. To dose the MCF-7, HCC1954, MDA-MB-231, and MDA-MB-468 cells with DNA-SWCNTs, the culture media were removed, cells were washed twice with PBS (2x, 1 mL per wash), and introduced to 2 mg L^−1^ DNA-SWCNTs diluted in the corresponding cell culture media, then incubated for 60 min to allow for cellular internalization. The SWCNT-containing media was then removed, the cells were washed twice with PBS (2x, 1 mL per wash), followed by incubation with 500 μL of trypsin at 37 °C and 5% CO_2_ for 7 min to facilitate cell detachment. Cells were then resuspended in 2 mL of complete growth medium and reseeded into a new culture plate. Then, dishes were incubated for 6 h before imaging. To dose the MCF10A with DNA-SWCNTs, the culture medium was removed, and the cells were rinsed with DPBS (2x, 1 mL per wash) followed by incubation with 500 μL of trypsin–EDTA at 37 °C and 5% CO_2_ for 15 min to detach the cells. Trypsin activity was neutralized with 500 μL of 0.1% soybean trypsin inhibitor in DPBS, and the cell suspension was subsequently centrifuged (400 × g, 10 min). The resulting cell pellet was resuspended in 2 ml of complete growth medium, reseeded into a new culture plate, and incubated for 6 h before imaging. NIR fluorescence spectra were acquired after 6 hours using a fiber optics probe spectroscopy system described previously. Acquired fluorescence was processed using custom MATLAB scripts to eliminate background interference prior to spectral analysis.

### Confocal Raman Microscopy

Raman microscopy measurements were acquired using a WiTec Alpha300R confocal Raman microscope (WiTec, Germany) equipped with an LD EC Epiplan-Neofluar DIC 50x air objective, a 785 nm (1.58 eV) excitation laser (35 mW output measured at the sample), and a UHTS 300 spectrograph with a 300 lines/mm grating coupled to an Andor DR32400 CCD detector (−61 °C, 1650 × 200 pixels. All cell lines were exposed to 2 mg L^-1^ DNA-SWCNTs for 60 min, following the same protocol used for near-infrared fluorescence imaging and maintained in fresh culture medium for an additional 6 h before analysis. After incubation, cells were fixed with 4% paraformaldehyde (PFA Electron Microscopy Sciences) in PBS for 15 min prior to imaging. Following fixation, cells were rinsed three times with PBS (3x, 2 mL per wash) and maintained in PBS before imaging. To minimize spectral interference during Raman measurements, PBS was removed, and the cells were rinsed with deionized water. The samples were then air-dried, and Raman spectra were acquired after complete drying.^52^ Individual cell regions were scanned, and Raman spectra were collected at 1.2 × 1.0 μm spatial intervals, with a 0.6 s integration time per spectrum, to generate hyperspectral single-cell Raman images. Raman spectra were acquired from more than 10 individual cells per cell line, and preprocessed prior to analysis to reduce instrumental noise and background contributions. The Savitzky–Golay (S–G) smoothing and cosmic-ray rejection (CRR) algorithms were used to improve the signal-to-noise ratio, and polynomial subtraction was used to eliminate fluorescence background and spectral drift in the WiTec Project 5.2 software prior to analysis. Processed hyperspectral datasets were further analyzed using customized MATLAB scripts to extract spectral band intensities, including integrated G-band intensity and RBM band intensities for each individual cell. The cell areas were quantified using image analysis. The Raman spectrum from each cell was normalized to its corresponding cell area and subsequently averaged for each cell line. Peak fitting was then performed independently using Gaussian functions to quantitatively compare SWCNT-related Raman features across different cell types.

### Cell Viability Assay

Cell viability following DNA-SWCNT exposure was evaluated using Annexin V and propidium iodide (PI) staining. Cells were plated on 35mm tissue-culture dishes (CellTreat) and cultured overnight at an initial seeding density of 3 × 10^5^ cells/cm^2^. After 24 hours, the culture media was replaced with 2 mg-L^-1^ DNA-SWCNTs diluted in the corresponding growth media, incubated for one hour, re-plated into the new petri dish, and incubated for an additional 6 hours. Following incubation, cells were collected and stained using the Dead Cell Apoptosis Kit V13242 (Invitrogen) following the manufacturer’s instructions. Fluorescence images of stained cells were acquired using a Cellometer Vision CBA image cytometer (Nexcelom Bioscience), and datasets were analyzed using customized MATLAB scripts. For each cell line, fluorescence gating thresholds for Annexin V and PI positivity were established using DNA-SWCNT-free negative-control cells and then applied consistently to the corresponding DNA-SWCNT-exposed samples.

### Data Collection for Machine Learning

Near-infrared spectral data (1100-1250 nm) from individual cells were collected for machine-learning classification of healthy and cancerous breast cell types. Each cell type had 600 data points. 3000 data points were used for training, and 600 for independent model testing. Each data point represents the fluorescence spectrum of an individual cell, which contains 15 features used in the model.

### Data Processing for Machine Learning

Cell borders in white-light microscopy images were segmented using Cellpose, a generalist deep-learning-based algorithm for cellular segmentation. The analysis was performed on Ubuntu 24.04.2 LTS using a Python-based Cellpose workflow. The resulting regions of interest (ROIs) were applied to the corresponding hyperspectral fluorescence datasets and processed using a custom MATLAB graphical user interface (GUI) to extract single-cell spectral features, including the center wavelength, peak intensity, and full width at half maximum (FWHM) of each emission band, as well as the broadband fluorescence intensity for each cell. All extracted spectral features were normalized using z-score standardization to reduce variability across samples. These features were used to train a machine learning classifier.

### Machine Learning Classification

The features were fed into an ensemble model for classification using the Classification Learner app in the MATLAB 2024b App Toolbox. Fifteen predictors, including center wavelength, normalized intensity, cell area, full width at half maximum (FWHM), and band ratio values, were used to train the model. 80% of the data set was used for the 10-fold cross-validation method to train the model. A stratified 20% holdout dataset is used as an independent test dataset. Model hyperparameters were optimized using Bayesian optimization with 100 iterations to enhance classification performance. Model performance was evaluated using classification accuracy, sensitivity, specificity, precision, and F1-score standards.

### Sample Preparation for Recapitulation Experiments

We first recapitulate the constant concentration of 5 mg L^-1^ DNA-SWCNTs in solutions with different intercellular conditions, collect their NIR fluorescence spectra immediately, and normalize them to the maximum for radiometric analysis. Peak fitting was then performed on the Band 2 fluorescence spectra using a Gaussian function to plot the normalized peak-2 intensity as a function of pH, hydrogen peroxide concentration, and ascorbic acid concentration. Finally, the variation was fitted with a rational function in OriginPro 2024b.

### Statistical Analysis

Statistical analyses were performed using OriginPro 2024b and MATLAB software. Due to the non-normal distribution and heterogeneity commonly observed in spectral datasets, statistical significance between experimental groups was evaluated using the non-parametric Kruskal–Wallis analysis of variance (ANOVA), followed by Dunn’s multiple comparisons test for pairwise group comparisons. A significance level of p<0.05 was considered statistically significant.

## Results and discussion

To create NIR fluorescent and biocompatible nanosensors, HiPco SWCNTs were first functionalized with single-stranded DNA oligonucleotides via probe-tip sonication and ultracentrifugation. In agreement with our previous studies^31^, the resulting DNA-SWCNT solution is highly purified from residual catalyst and other impurities as evidenced by high peak-to-valley ratios in absorbance scans, strong Raman signatures, and bright NIR fluorescence **(Figure S1)**. The DNA sequence (GT)_6_ was chosen over other sequences since we have shown that the relatively short DNA strands exhibit lower intracellular stability,^25,31,53^ enabling easier displacement by amphiphilic molecules and thereby enhancing chirality-dependent fluorescence modulation.

We next investigated the uptake and biocompatibility of the DNA-SWCNTs within breast cancer cells. Five distinct cell lines of MCF10A, MCF7, HCC1954, MDA-MB-231, and MDA-MB-468 were utilized as models of non-tumorigenic control and the luminal, HER2+, and triple-negative breast cancer cells^54^, respectively (**Table S1**). It is expected that DNA-SWCNTs enter into the breast cancer cells via endocytosis and localize to the lysosomes^55^. Confocal Raman microscopy was used to confirm the uptake of DNA-SWCNTs in each cell line.^29,56,57^ The G-band, originating from the sp²-hybridized carbon lattice of SWCNTs^58^, exhibits a linear correlation with nanotube concentration^59^ and was therefore used to quantify intracellular SWCNT uptake. **Figure 1a** presents transmitted-light images, Raman-integrated G-band intensity maps, and corresponding overlay images of the same individual cells across all cell lines, confirming efficient uptake of DNA-SWCNTs in all cell types. The quantitative results of the G-band are summarized in **Figure 1c**. Among all cell lines, MDA-MB-468 exhibited significantly higher SWCNT uptake than all the other cell types. Previous studies have reported that Epidermal Growth Factor Receptor expression on the surface of MDA-MB-468 cells is exceptionally high (∼1.3×10^6^ receptors per cell)^21^. Other studies have further demonstrated that cellular uptake of targeted nanocarriers positively correlates with this receptor density^22^, which may explain the enhanced uptake of SWCNTs observed in this cell line. The HCC1954 and MCF7 cell lines also exhibited significantly different uptake levels. However, other cell lines did not show significantly different from each other. The average G-band intensities normalized to cell area for MCF10A, MCF7, HCC1954, MDA-MB-231, and MDA-MB-468 were 1.126, 2.093, 0.714, 0.942, and 3.742, respectively.

**Figure 1.**
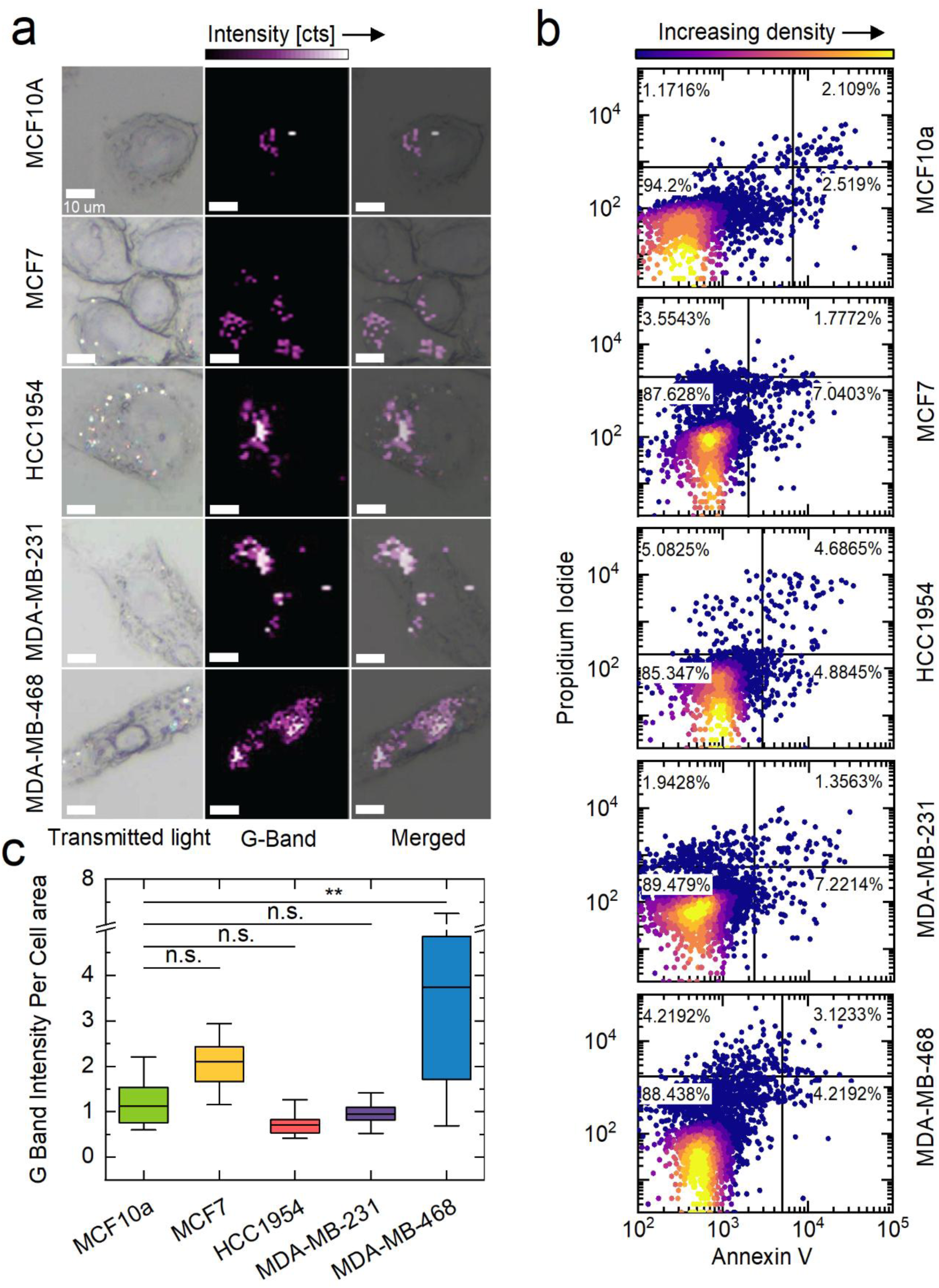
Confocal Raman microscopy characterization and cell viability assay for all breast cancer cell lines and a healthy breast control cell line. (a) Transmitted light images, magnified Raman G-band images, and merged images at 50x, separated by cell type. (b) Scatter plots for the Annexin V/PI viability assay following continuous culture in a 2 mg-L⁻¹ DNA-SWCNTs solution for 6 hours. Gatings are determined from negative controls. Percentages of cells in each quadrant are shown. (c) Box plot showing the G-band intensity per cell area. Whiskers extend to the minimum and maximum values (n ≥ 10 cells per condition). Statistical significance was assessed using a Kruskal–Wallis ANOVA followed by Dunn’s multiple comparisons test (**p<0.01).

Following the confirmation of cell uptake, the cytotoxicity of DNA-SWCNTs was evaluated as a function of cell type using an apoptosis assay based on Annexin V and propidium iodide (PI) staining.^60^ Shown in **Figure 1b and S2** are the PI vs Annexin V intensity scatter plots. The bottom-left quadrant corresponds to viable cells, the bottom-right quadrant to early apoptotic cells, the top-right quadrant to late apoptotic cells, and the top-left quadrant to necrotic cells. The DNA-SWCNTs at the chosen incubation concentration of 2 mg L^-1^ does not induce significant apoptosis or necrosis in any of the breast cancer or healthy control cell lines.

Near-infrared hyperspectral fluorescence microscopy was performed on all 5 cell types with internalized DNA-SWCNTs. To identify the optimal timepoint for fluorescence imaging in this study, MCF7 cells were investigated at 4, 6, or 24 hours after an initial incubation period with DNA-SWCNTs **(Figure S3)**. The fluorescence signal reached a maximum at 6 hrs post-incubation, in agreement with our previous study^53^. We therefore selected the 6 hour incubation timepoint for all subsequent fluorescence experiments. **Figure 2a** presents transmitted-light, broadband NIR fluorescence, and overlay images for all cell types 6 hours after dosing. Bright NIR fluorescence is observed in all 5 cell types with notably bright signal found in MDA-MB-468 cells. Also, cell areas were quantified via image analysis as described in **Figure S5f**. **Figure 2b** presents the average broadband NIR fluorescence intensity for each cell type normalized by the corresponding cell area (n=200). As predicted from the images, the area-normalized broadband intensity is highest in MDA-MB-468. In contrast, MCF10A exhibits the lowest normalized intensity, followed by HCC1954, consistent with its larger average cell area, which reduces the normalized intensity per unit area. We examined two distinct bright emission bands in the range 1100-1250 nm that we term Band 1 and Band 2 which are dominated by the (9,4) and (8,6) SWCNT^31^. **Figure 2c** presents the average fluorescence spectra of n=200 cells across all 5 cell lines at the 6 hour time point, normalized to the corresponding Band 1 peak intensity. Distinct variations in normalized intensities, peak center wavelengths, and full widths at half maximum (FWHMs) are observed among the evaluated cell lines. All breast cancer cell lines demonstrated distinct modulation relative to healthy MCF10A cells in both bands. Quantitative analysis on the collected spectra were performed by peak fitting and examining the spectral features. **Figure 2d** presents a box plot comparing the Band 2 center wavelength across all cell types. All breast cancer cell lines exhibited significantly red-shifted fluorescence emission compared to the healthy MCF10A cells. Specifically, MCF-7, HCC1954, MDA-MB-231, and MDA-MB-468 exhibited red shifts of 4.3, 13.5, 10.4, and 14.1 nm in the Band 2 center wavelength, respectively, relative to MCF10A. This finding is our first indication that DNA-SWCNTs can be used as NIR fluorescence sensors for the in vitro detection of breast cancer.

**Figure 2.**
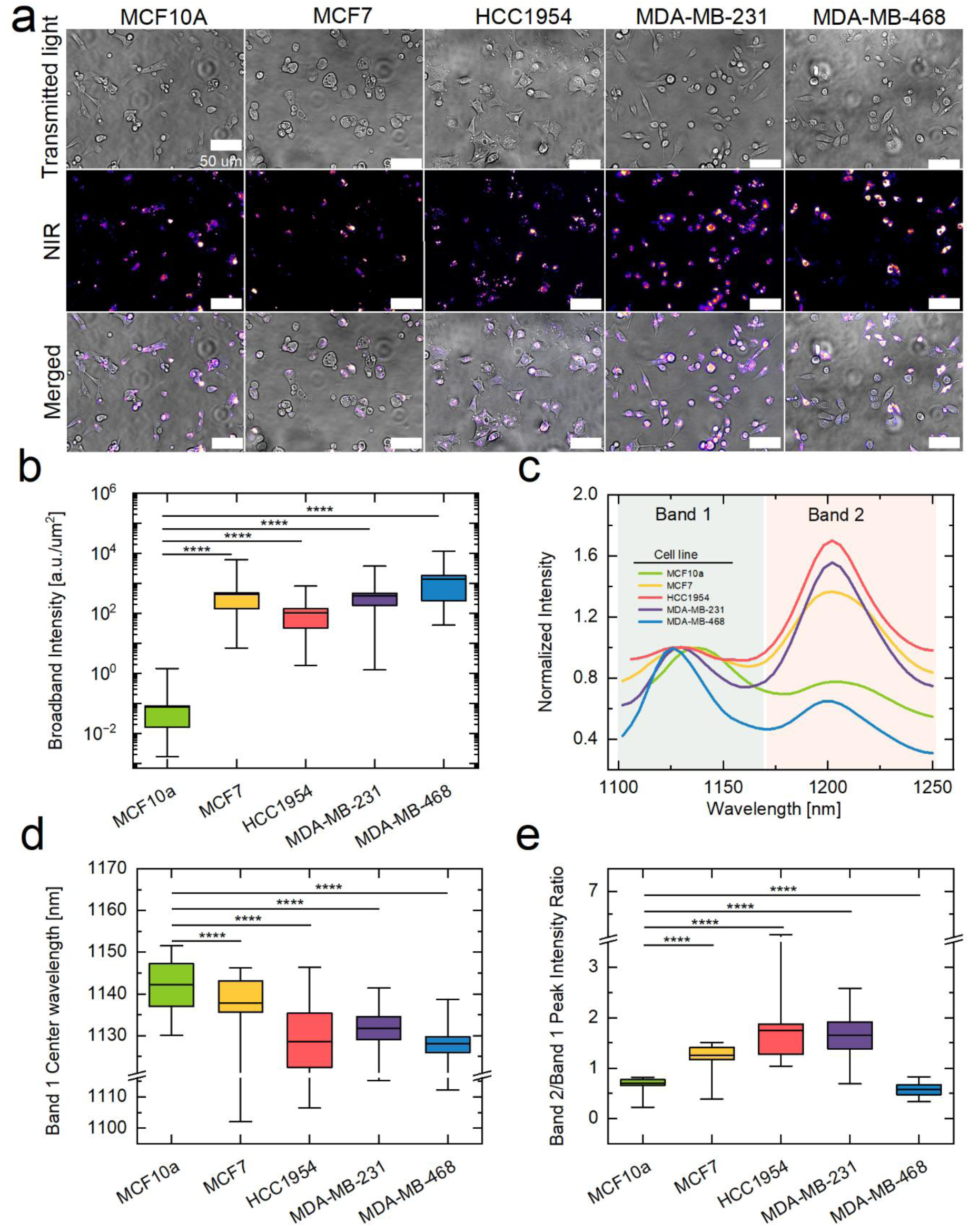
NIR fluorescence hyperspectral microscopy and spectral feature analysis of DNA-SWCNTs in healthy and cancerous breast cell types 6 hours after internalization. (a) Transmitted light, broadband NIR fluorescence (900–1600 nm), and merged images for all examined cell types. (b) Box plot of average broadband intensity per cell area across all cell types. (c) Average NIR spectrum of DNA-SWCNTs within each cell type, with Band 1 peak intensity normalized to 1. (d) Box plot of Band 2 center wavelength. (e) Box plot comparing Band 2/Band 1 peak intensity ratios across all cell types. Boxes represent 25–75% of the data, horizontal lines denote the mean, and Whiskers extend to the minimum and maximum values (n = 200 cells per condition). Statistical significance was assessed using a Kruskal–Wallis ANOVA followed by Dunn’s multiple comparisons test (****p < 0.0001).

As another quantitative measure of variation among the cell types, **Figure 2e** presents a comparison of the peak-intensity ratios of Band 2 versus Band 1. Statistically significant differences between all cell types are observed. MCF10A and MDA-MB-468 showed lower Band 2 intensities than their corresponding Band 1 intensities, giving Band 2/Band 1 intensity ratios of 0.703 and 0.575, respectively. However, MCF7, HCC1954, and MDA-MB-231 displayed higher Band 2 than Band 1 peak intensity ratios of 1.258, 1.745, and 1.658, respectively. Broadly, we attribute the observed variations in NIR fluorescence spectra to differences in the endosomal microenvironment among the cell types, which can include factors such as pH, ion concentrations, degradative enzymes, etc^61–63^, which can be resolved in future studies

In order to discriminate cancerous from healthy breast cells, machine-learning analyses were performed to evaluate whether subtle spectral differences across multiple correlated features could collectively distinguish the cell populations. As an initial unsupervised approach, principal component analysis (PCA) was used to visualize the overall variance and clustering behavior of the extracted spectral features. The results of the corresponding analysis of the remaining spectral features used for machine-learning classification are presented in **Figure S5**. The PCA score plot (**Figure S7**) shows substantial overlap between the non-cancerous and cancerous populations, indicating that the variations are complex and not linearly separable.^64^ This complexity necessitates the use of supervised machine learning models for robust and accurate classification^64^ of non-cancerous and cancerous cells and for sub-typing them based on NIR fluorescence features of the intracellular SWCNTs. All features were normalized using z-score standardization to reduce the influence of data variability^65^. Additionally, highly correlated features were excluded because redundant variables contribute limited new information and can introduce noise into the model. Although 20 features were initially extracted, 15 NIR fluorescence features were ultimately selected using a Pearson correlation coefficient threshold (|r| < 0.75) to maintain low inter-feature redundancy. The results of the Pearson correlation coefficient of selected features are shown in **Figure 3a and Figure S8**. Notably, Band 1 intensity exhibited strong correlations (|r| > 0.75) with multiple features and was therefore excluded from the machine learning model.

**Figure 3.**
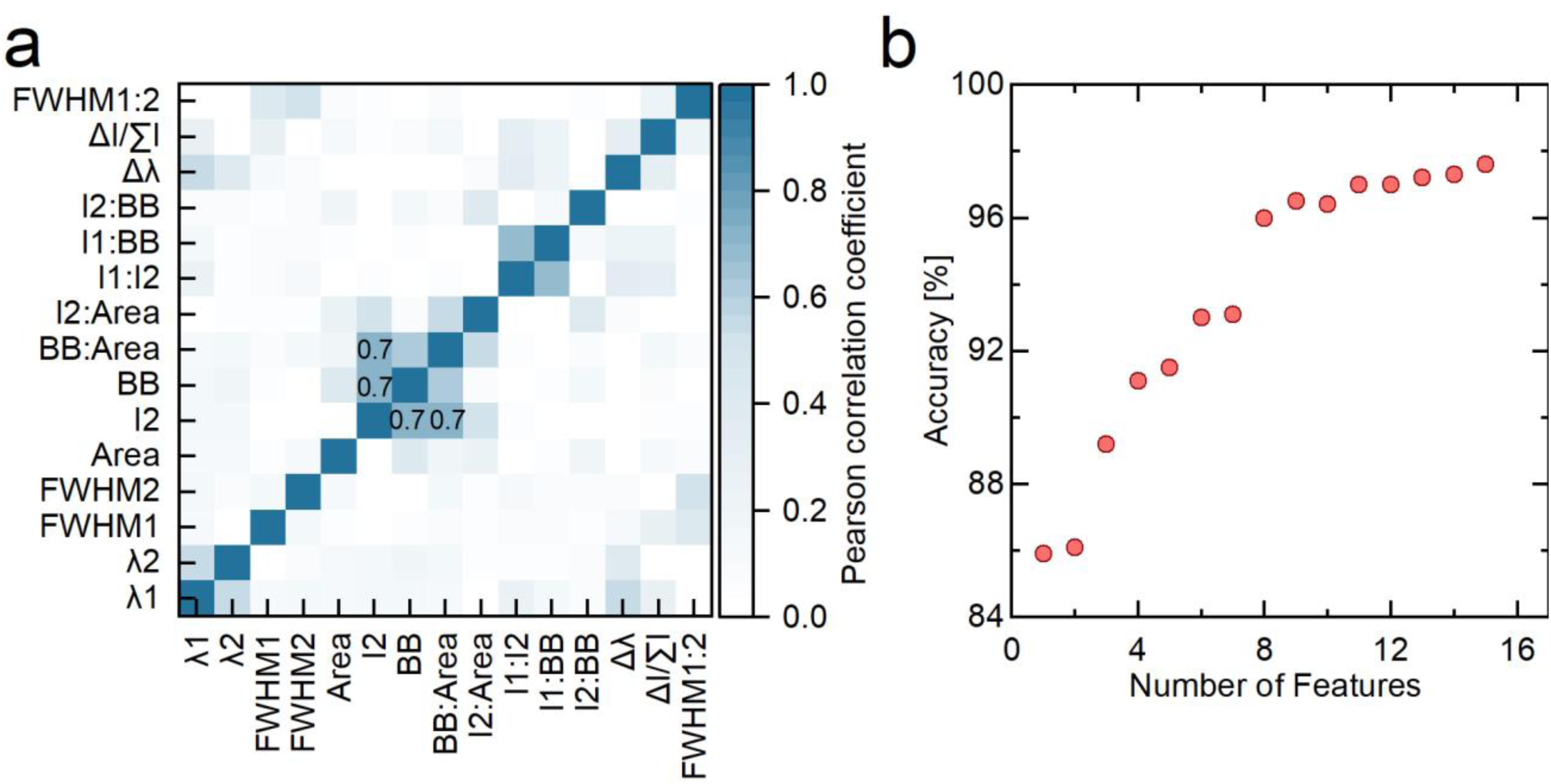
Feature engineering for breast cancer cell type discrimination. (a) The heat map shows the Pearson correlation coefficients among all features used in the machine learning model. Features with r > 0.75 were removed. (b) Breast cancer detection model performance as a function of the increasing number of sensor features.

To determine whether the NIR fluorescence spectra can differentiate healthy vs. cancerous cells, an optimizable ensemble algorithm was applied to the NIR fluorescence features of 600 individual MCF10A cells and 150 individual cells from all four breast cancer cell lines (600 cells total). First, the Kruskal–Wallis ANOVA test^66^ was performed to identify the most significant spectral features, as the data are not normally distributed (**Figure S9a**). An optimized AdaBoost ensemble^49^ model with decision tree learners was then implemented to evaluate classification performance as a function of the number of features, ranked from most to least important (**Figure 3b**). In order to tune the machine learning model hyperparameters, Bayesian optimization was performed over 100 iterations. The results are summarized in **Table S3**. A 10-fold cross-validation approach was employed to train the model. The optimized model was applied to 20% of the dataset as an independent test set.

The binary classification results for breast cancer detection are presented in **Figure 4**. The model achieved accuracies of 97.52% and 97.68% on the validation and test sets, respectively, demonstrating that it generalizes well to unseen data and exhibits minimal overfitting (**Figure 4a**). The receiver operating characteristic (ROC) curves for both cancerous and non-cancerous classes in the breast cancer detection model are shown in **Figure 4b**. The corresponding area under the curve (AUC) for cancerous and non-cancerous classes are 99.48% and 99.47%, respectively. **Figure 4c** exhibits the validation and test accuracies for each class. The results showed slightly higher classification performance for cancerous cells than for non-cancerous cells in both datasets. In the validation (and test) sets, cancerous cells achieved classification accuracies of 97.74% (97.92%), compared to 97.32% (97.54%) for non-cancerous cells. The trend further confirmed the model’s minimal bias and strong generalization. Moreover, precision, which reflects the fraction of positive predictions that are correct; sensitivity, which measures the ability of the model to correctly identify true positive cases; specificity, which evaluates the ability to correctly identify true negative cases; and the F1 score, which represents the harmonic mean of precision and sensitivity, were assessed to comprehensively evaluate the classification performance of the detection model.^52^ The results of the class-specific matrix are summarized in **Figure 4d**. Precision and specificity were higher for the cancerous class (97.76% and 97.76%, respectively) compared to the non-cancerous class (97.34% and 97.32%, respectively). In contrast, sensitivity was slightly higher for the non-cancerous class (97.76%) than for the cancerous class (97.32%). Consequently, the F1-score was identical for both classes (97.54%). These results support the effectiveness of the proposed spectral fingerprinting of SWCNTs for detecting breast cancer cells versus a healthy control at the single-cell level.

**Figure 4.**
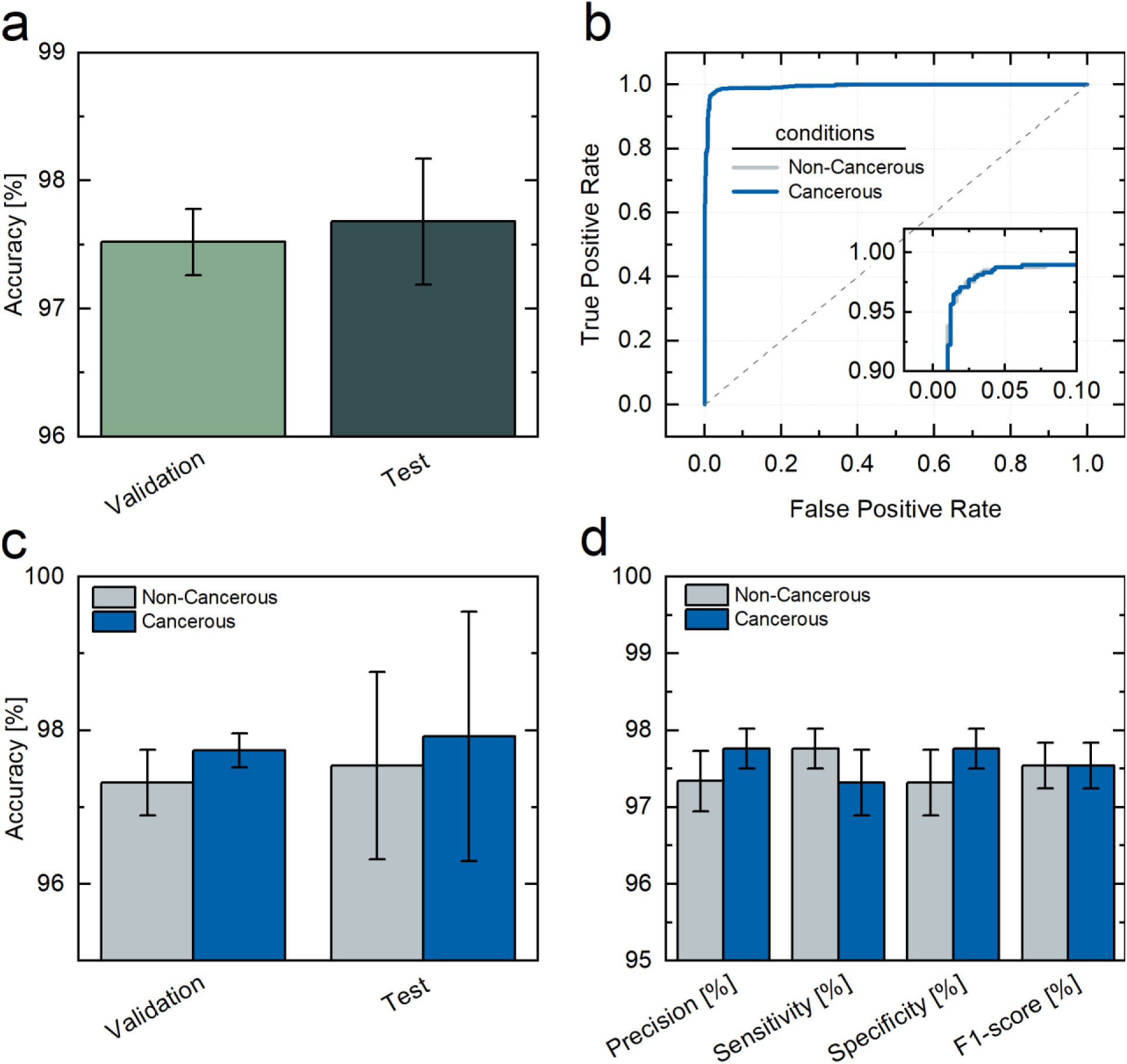
Machine learning classification for the detection of cancerous cells. (a) Bar graph indicating the accuracy of validation and test data. (b) ROC curve for cancerous and non-cancerous cells, indicating true positive and false negative rates. (c) Bar graph indicating the accuracy of validation and test data across cancerous and non-cancerous cell clusters. (d) Bar graph showing precision, sensitivity, specificity, and F1-score for cancerous and non-cancerous cells. Whiskers extend to the minimum and maximum values (n = 5).

After achieving remarkable results in breast cancer versus healthy cell *in vitro* detection, we assess whether NIR fluorescence spectra from internalized DNA-SWCNTs can distinguish between breast cancer cell line subtypes by applying another ensemble algorithm to their fluorescence spectral features. Bayesian optimization (**Table S4**) over 100 iterations, and a 10-fold cross-validation with 20% of the dataset as an independent test set was employed for the NIR fluorescence features of 600 individual cells from the 5 cell lines. The results of classification performance as a function of the number of features are shown in **Figure S9b**, and the classification results for cell typing are presented in **Figure 5**. **Figure 5a** compares classification performance on the validation and test datasets, with accuracies of 95.1% and 95.43%, respectively. **Figure 5b** shows ROC curves for all breast cancer and healthy cell types with AUC values of 99.94%, 99.75%, 99.03%, 99.03%, 99.77% corresponding to MCF10A, MCF7, HCC1954, MDA-MB-231, and MDA-MB-468 cell types, respectively. **Figure 5c** presents the accuracies for all classes in both the validation and test datasets, with performance ordered from highest to lowest across HCC1954, MCF10A, MDA-MB-468, MCF7, and MDA-MB-231. The corresponding validation and test accuracies are shown in **Table 1**. These results support minimal model bias and robust generalization of breast cancer cell typing.

**Figure 5.**
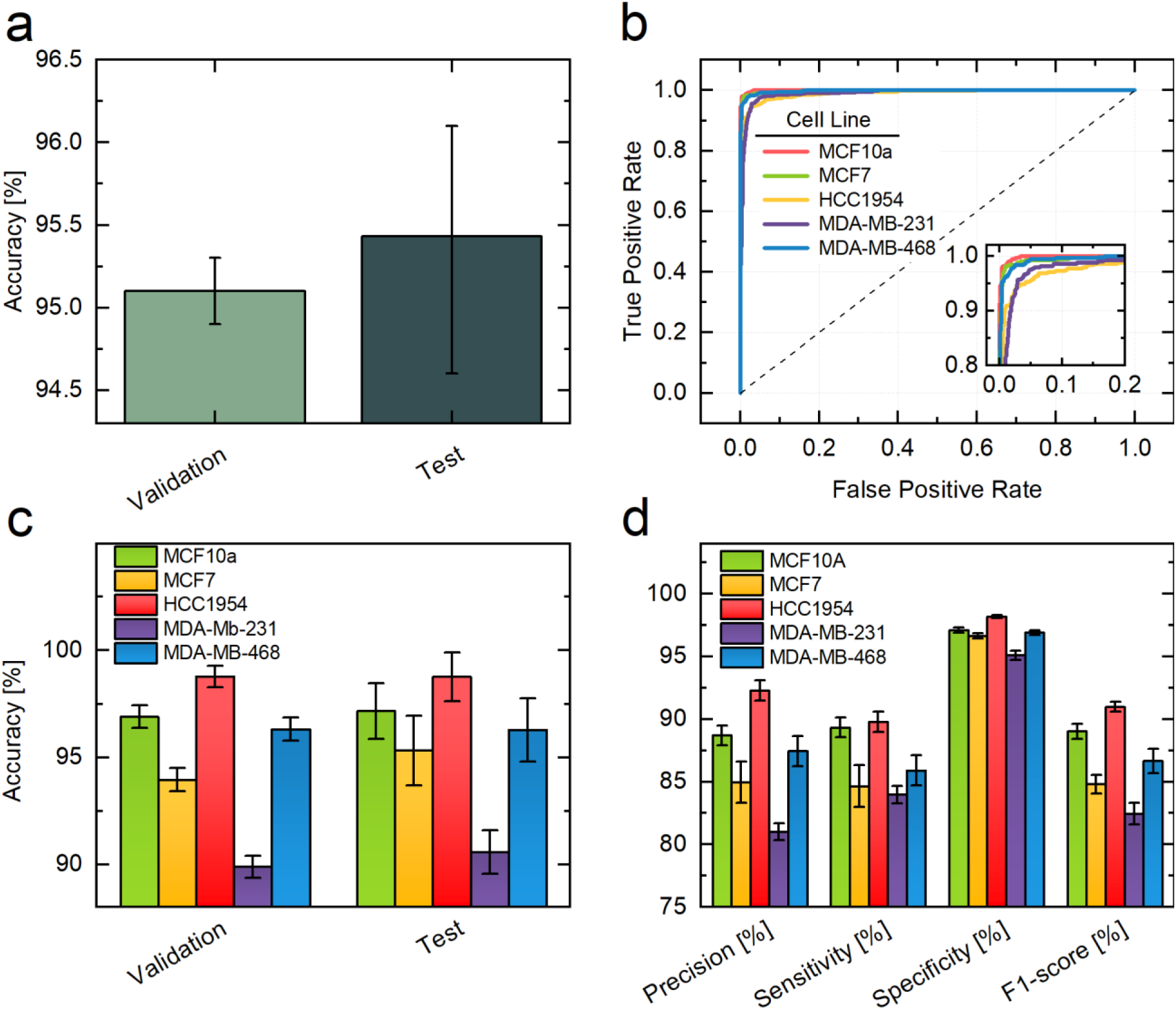
Machine learning classification for the cell typing of breast cancer. (a) Bar graph indicating the accuracy of validation and test data. (b) ROC curve for different breast cancer cell types and a healthy breast control cell line, indicating true positive and false negative rates. (c) Bar graph indicating the accuracy of validation and test data across all cancer cell types and a healthy breast control cell line. (d) Bar graph showing precision, sensitivity, specificity, and F1-score for breast cancer cell types and a healthy breast control cell line. Whiskers extend to the minimum and maximum values (n = 5).

**Table 1.** Performance of the model for breast cancer cell type discrimination.

| Cell line | validation accuracy (%) | Test accuracy (%) | Precision (%) | Specificity (%) | Sensitivity (%) | F1-Score (%) |
| --- | --- | --- | --- | --- | --- | --- |
| MCF10A | 96.9 | 97.17 | 88.70 | 97.12 | 89.32 | 89.02 |
| MCF7 | 93.95 | 95.32 | 84.96 | 96.66 | 84.64 | 84.80 |
| HCC1954 | 98.87 | 98.77 | 92.26 | 98.18 | 89.76 | 90.98 |
| MDA-MB-231 | 89.88 | 90.58 | 81.02 | 95.10 | 83.96 | 82.46 |
| MDA-MB-468 | 96.32 | 96.28 | 87.44 | 96.9 | 85.90 | 86.66 |

Precision, sensitivity, and specificity of the cell typing machine learning model were also assessed. The class-specific metrics are shown in **Figure 5d**. The HCC1954 class achieved the highest, and the MDA-MB-231 cell category the lowest precision, sensitivity, and specificity. Specificity showed the highest values, while precision and sensitivity exhibited moderate variation among cell lines. The results are summarized in Table 1. Collectively, these results demonstrate that the SWCNT spectral fingerprinting platform has potential for discrimination among distinct breast cancer cell types with high classification performance. Although this was a small screen as a proof-of-concept, with 1 cell line representing each of luminal and HER2+, and two cell lines representing TNBC, spectral fingerprinting of SWCNTs coupled with a machine learning approach shows that we can not only distinguish between cancer vs. non-cancer, but also potentially between each breast cancer cell line. Although we do not yet know the mechanism by which SWCNT discriminates between them, this is a promising potential method to detect BC subtypes and support clinical decision making. In future studies, we propose a larger screen with many cells representing each subtype and discernment for how the subtype influences the SWCNT fingerprint.

After successfully detecting breast cancer cell types in vitro, we aimed to identify the reasons for the observed changes in their fluorescence spectra. We identify two primary factors that may account for differences in broadband intracellular fluorescence intensity of DNA-SWCNTs between cell lines: (1) variations in total DNA-SWCNT uptake and (2) fluorescence quenching induced by SWCNT aggregation within intracellular compartments.^31^ To analyze the uptake and degree of aggregation of SWCNT, characteristic Raman features of SWCNTs, including the G-band and radial breathing mode (RBM), were studied. The results of Raman spectroscopy are shown in **Figure 6**. The most informative regions of the average Raman spectra, normalized by cell area for each cell line, are presented in **Figure 6a**. Additionally, Recall from **Figure 1c** that variations in DNA-SWCNT uptake were indeed observed between cell types, as indicated from the G-band of the SWCNT Raman spectra. Plotting broadband NIR fluorescence intensity versus G-band Raman signal for each cell type, clustering is observed between the cancer cell types, leaving the healthy MCF10A control cell type quantitatively distinct (**Figure 6b**).^31,58,59,67,68^ To examine the possibility of fluorescence quenching due to SWCNT aggregation, Raman microscopy focusing on the radial breathing mode (RBM) of SWCNTs was performed on healthy and cancerous breast cell lines containing the internalized DNA-SWCNTs.

**Figure 6.**
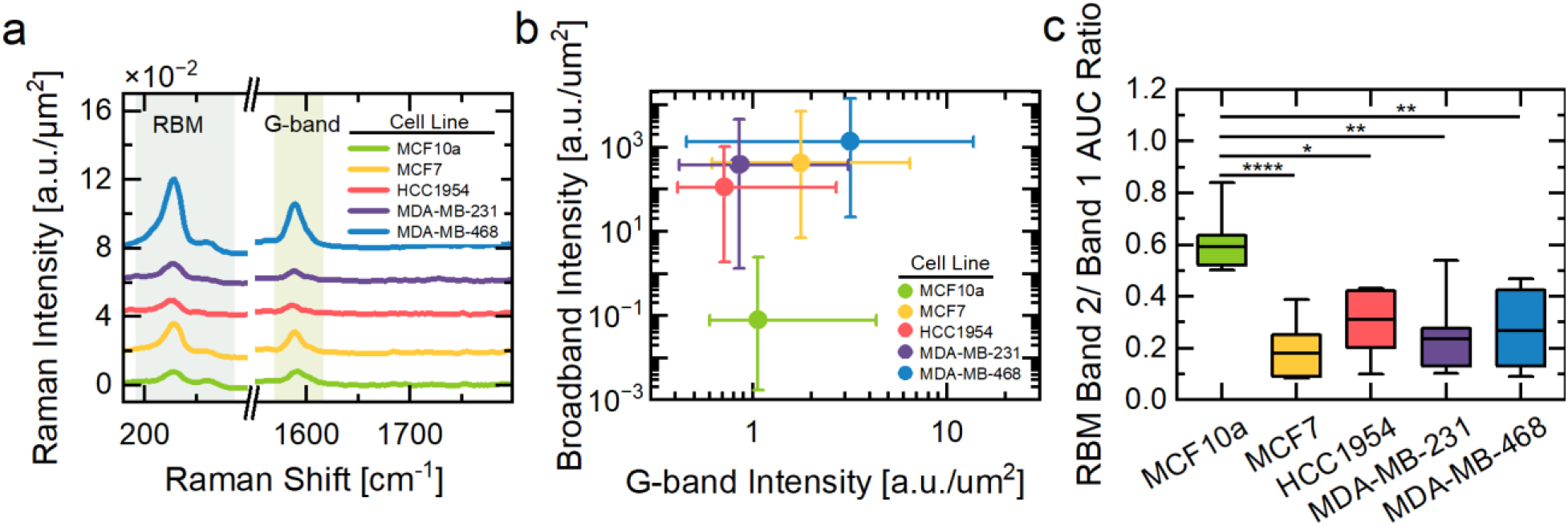
Confocal Raman microscopy analysis of SWCNT uptake by different cell types. (a) Stacked line plot showing the spectrum of average normalized Raman intensity per cell area. Spectra of MCF7, HCC1954, MDA-MB-231, and MDA-MB-468 are shifted by 2, 4, 6, and 8, respectively, for visual clarity. (b) Scatter plot showing the broadband intensity per cell area versus G-band intensity per cell area across different cell types. Whiskers extend to the minimum and maximum values (n ≥ 10 cells per condition). (c) Box plot showing the ratio of Band 2/Band 1 AUC of the radial breathing mode (RBM) in the Raman spectrum of each cell type. Boxes represent 25–75% of the data, horizontal lines denote the mean, and whiskers extend to the minimum and maximum values (n ≥ 10 cells per condition). Statistical significance was assessed using a Kruskal–Wallis ANOVA followed by Dunn’s multiple comparisons test (*p<0.05, **p<0.01, ****p < 0.0001).

The RBM region of the intracellular DNA-SWCNTs exhibited two peaks that we named RBM Band 1, ranging from 217 to 242 cm^-1^, corresponding to the (10,5), (11,3), and (12,1) SWCNT chiralities, and RBM Band 2, located between 250 and 275 cm^-1^, which is associated with the (9,4) and (10,2) chiralities.^31^ E_22_ transition energies of the RBM Band 1 chiralities are closely matched to the laser excitation energy (1.58 eV), placing these nanotubes under resonant excitation conditions and therefore producing a stronger RBM Band 1 intensity relative to Band 2. However, upon intracellular aggregation, the E_22_ transition energies undergo red shifting due to inter-nanotube electronic coupling and changes in the local dielectric environment. As a result, the chiralities associated with RBM Band 2 shift closer to resonance with the 1.58 eV excitation source, resulting in stronger RBM Band 2 intensity, while the RBM Band 1 chiralities progressively move out of resonance. Consequently, the ratio of the areas under the RBM Band 2 and RBM Band 1 curves is attributed to intracellular aggregation of DNA-SWCNTs, consistent with previous reports.^31,69^ **Figure 6c** presents the RBM Band 2/Band 1 area under the curve (AUC) ratio for all cell types. MCF10A exhibited the statistically highest aggregation level relative to all examined cell types. However, other cell lines did not exhibit significantly different degrees of aggregation. The average RBM Band 2/Band 1 AUC ratios for MCF10A, MCF7, HCC1954, MDA-MB-231, and MDA-MB-468 were 0.59, 0.18, 0.31, 0.24, and 0.27, respectively.

Overall, Raman spectra showed that MDA-MB-468 exhibits the highest G-band intensity and, correspondingly, the strongest broadband fluorescence, confirming that its enhanced optical response is attributable to SWCNT uptake. In contrast, MCF10A displays relatively low broadband fluorescence intensity despite approximately the same SWCNT uptake as the other cell lines except MDA-MB-468, consistent with its high intracellular aggregation observed from the RBM analysis, confirming that SWCNT fluorescence is quenched in this cell line due to aggregation.

We hypothesize that the observed changes in fluorescence spectra, including shifts in center wavelength and changes in intensity ratios among different chiral species, are attributable to variations in the lysosomal environment of different breast cancer cell phenotypes.^25^ Previous studies have demonstrated that the endosomal microenvironment differs substantially among breast cancer cell types in terms of redox activity, pH, ionic composition, metabolite concentration, and biomolecular content.^61–63^ Together, these variations modulate the DNA-SWCNT fluorescence spectrum.^25^ Furthermore, modulations in the fluorescence spectra of intracellular DNA-SWCNTs generally cannot be attributed to individual biomarkers. To confirm the hypothesis, we focus on the broader influence of intracellular redox and oxidative microenvironments, pH, ionic composition (K⁺, Ca²⁺, and Mg²⁺), and ionic strength arising from variations in NaCl concentration on the optical behavior of DNA-SWCNTs in the solution phase.

The results of the solution phase DNA-SWCNT recapitulation studies are shown in **Figure 7** and **Figure S11-S12. Figure 7a, d** represents the effect of varying pH, ranging from 4.5 to 8, on the fluorescence spectra of SWCNTs. The results reveal that the ratio of Band 2/Band 1 increases with increasing pH. A similar trend is observed in the case of an increasingly oxidative environment as demonstrated by the addition of hydrogen peroxide (**Figure 7b,e**), which is in agreement with a previous study.^70^ Finally, **Figure 7c,f** explores the effect of a reducing environment on solution-phase DNA-SWCNTs by addition of ascorbic acid. The Band 2/Band 1 ratio decreases with increasing ascorbic acid. We rationalize these findings by acknowledging that DNA-SWCNTs contained within the endosomal pathway of mammalian cells are subjected to characteristic variations in pH and redox as early endosomes progress to late endosomes and finally lysosomes.^30,31,55,70^ This pathway is highly cell type dependent, as evident in our NIR fluorescence data. Despite this complexity, our results are consistent with previous studies showing that MDA-MB-231 cells exhibit oxidative lysosomes and high autophagic flux. However, MDA-MB-468 cells exhibit a highly reducing environment, lower basal autophagy, and elevated dsDNA accumulation.^71–73^

**Figure 7.**
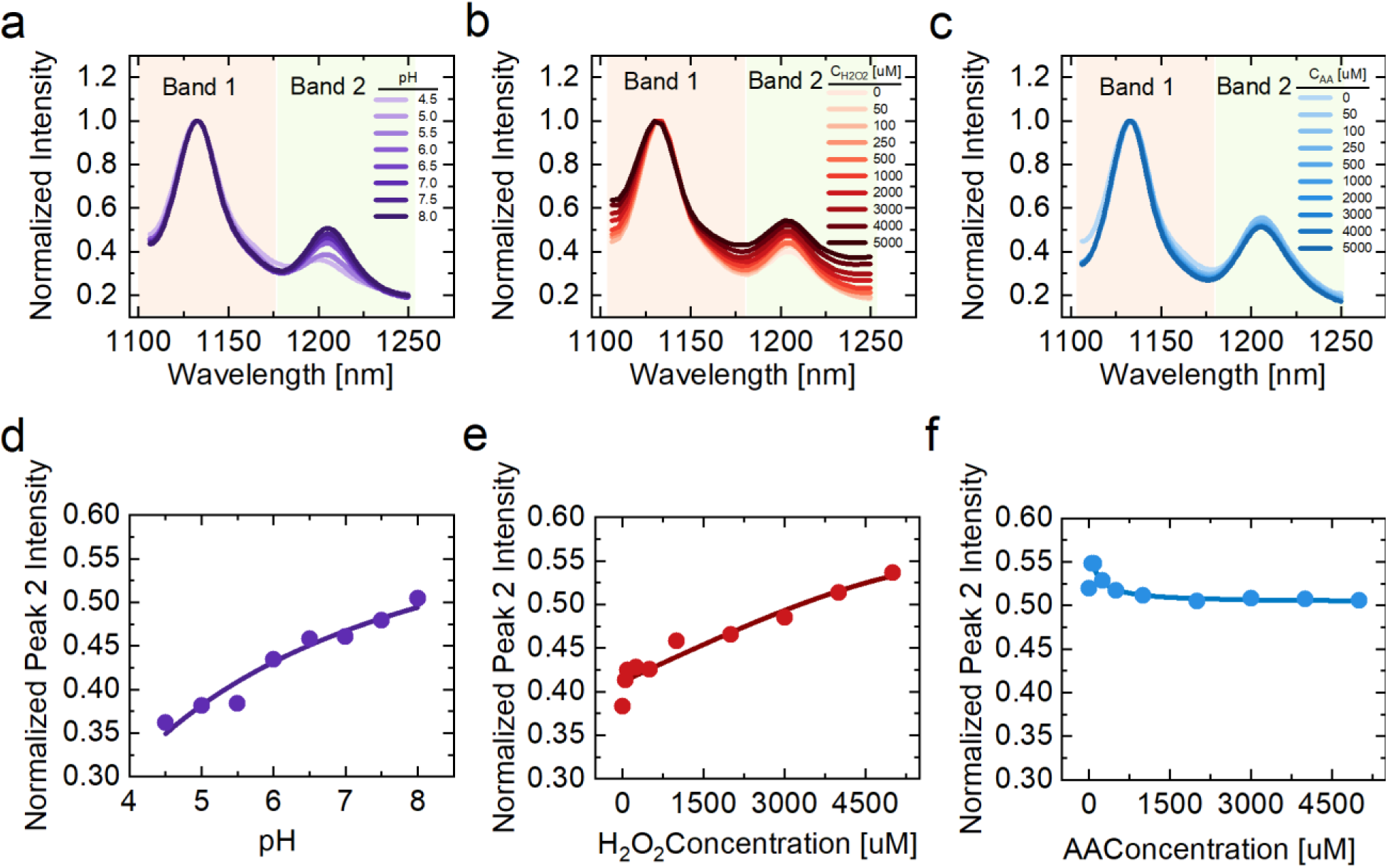
Solution-phase experiments recapitulating the NIR fluorescence response of the DNA-SWCNTs sensor to the endolysosomal environment. (a) Spectral response to varying pH environments, (b) varying hydrogen peroxide as an oxidation agent, and (c) varying ascorbic acid as a reducing agent. (d) Dose–response curve of normalized Peak 2 intensity as a function of environmental pH, (e) dose–response curve of normalized Peak 2 intensity as a function of hydrogen peroxide concentration, and (f) dose–response curve of normalized Peak 2 intensity as a function of ascorbic acid concentration.

Overall, Changes in pH, oxidative or reducing conditions produced measurable changes in the nanotube fluorescence response, demonstrating that DNA-SWCNT emission is sensitive to several physicochemical features known to vary across breast cancer cells. Therefore, the cell-type-dependent spectral patterns observed reflect the combined influence of these intracellular environmental parameters on DNA-SWCNT photophysics. This result further supports the use of DNA-SWCNTs as optical reporters of cell-dependent intracellular states and provides a mechanistic basis for their use in spectral fingerprinting of breast cancer cells.

## Conclusion

In this work, we demonstrated that DNA-SWCNT near-infrared fluorescence provides a high-accuracy spectral fingerprinting platform for detecting breast cancer and distinguishing breast cancer cell types at the single-cell level. After exposure to DNA-SWCNTs, each cell line produced reproducible nanotube emission features, including changes in band intensity, center wavelength, FWHM, and broadband fluorescence intensity. These spectral differences were sufficient to separate non-tumorigenic MCF-10A cells from breast cancer cells and to discriminate among molecularly distinct breast cancer cell models, including MCF7, HCC1954, MDA-MB-231, and MDA-MB-468.

The strong classification performance indicates that DNA-SWCNT optical responses capture cell-type-dependent differences in the cellular microenvironment rather than simple differences in nanotube exposure. Importantly, cell viability measurements showed that the nanotube exposure conditions used here did not induce significant apoptosis or necrosis, supporting the use of this approach for non-destructive cellular analysis. Moreover, Raman spectroscopy showed differences in nanotube uptake and aggregation across cell lines, with the highest SWCNT uptake observed in MDA-MB-468 and the strongest aggregation observed in MCF-10A. Fluorescence spectroscopy further revealed cell-type-specific emission profiles, including lower broadband intensity in MCF-10A, higher average Band 1 intensity in MDA-MB-468 and MCF-10A, and higher average Band 2 intensity in HCC1954, MCF7, and MDA-MB-231. When combined with an ensemble classification algorithm, these fluorescence-derived features enabled both breast cancer detection and breast cancer cell typing. Overall, this work establishes DNA-SWCNT spectral fingerprinting as a non-destructive optical platform for capturing cell-type-dependent differences and motivates its further development for rapid phenotypic profiling in breast cancer.

## Acknowledgment

This work was supported by the National Science Foundation (CAREER Award no. 1844536). TOC graphics and schematics were created using https://BioRender.com software.

## Notes

The authors declare no competing financial interest.

