## supporting information for "In Vitro Detection of Breast Cancer Cell Types Using Machine Learning-Assisted Spectral Fingerprinting of SWCNTs"

### Data Processing for Machine Learning

Cellpose was installed and executed on Ubuntu 24.04.2 LTS following a GitHub-hosted installation/debugging workflow for CellProfiler and Cellpose, with all image-segmentation parameters kept constant across the dataset.

**Table 1.** Breast epithelial and breast cancer cell models used for DNA-SWCNT spectral fingerprinting.

| Subtype | Cell Model | ER | PR | HER2 | Description |
|---|---|---|---|---|---|
| Healthy cell | MCF10A | | | | non-tumorigenic immortalized mammary epithelial cell line derived from fibrocystic breast tissue |
| Luminal | MCF7 | + | + | - | established from a pleural effusion in 1973, low invasive |
| HER2+ | HCC1954 | - | - | + | derived from a stage IIB primary tumor |
| Triple negative claudin low | MDA-MB-231 | - | - | - | highly invasive triple negative line with a claudin low, mesenchymal phenotype and high metastatic potential in xenograft models |
| Triple negative basal like | MDA-MB-468 | - | - | - | TNBC line belongs to the basal-like subtype, EGFR overexpressing |

**Table 2.** Spectral features extracted from single-cell DNA-SWCNT fluorescence images

| | Spectral Feature | Symbol |
|---|---|---|
| 1 | Center Wavelength of Band 1 | $\lambda_1$ |
| 2 | Center Wavelength of Band 2 | $\lambda_2$ |
| 3 | Broadband Intensity | $BB$ |
| 4 | Band 2 Intensity | $I_2$ |
| 5 | Full Width at Half Max of Band 1 | $FWHM_1$ |
| 6 | Full Width at Half Max of Band 2 | $FWHM_2$ |
| 7 | Cell Area | $Area$ |
| 8 | Broadband Intensity/ Cell Area | $\frac{BB}{Area}$ |
| 9 | Band 2 Intensity/ Cell Area | $\frac{I2}{Area}$ |
| 10 | Band 1 Intensity/ Band 2 Intensity | $\frac{I_2}{I_1}$ |
| 11 | Band 1 Intensity/ Broadband intensity | $\frac{I_1}{BB}$ |
| 12 | Band 2 Intensity/ Broadband intensity | $\frac{I_2}{BB}$ |
| 13 | Full Weight at Half Max of Band 1/Full Weight at Half Max of Band 2 | $\frac{FWHM_1}{FWHM_2}$ |
| 14 | Band Center Separation | $\lambda 2\text{-}\lambda 1$ |
| 15 | Band Intensity Contrast | $\frac{I2-I1}{I2+I1}$ |

**Table 3.** Optimized hyperparameters of the ensemble classification algorithm for breast cancer detection.

| Model Hyperparameters | range | Optimized Hyperparameters |
|---|---|---|
| Ensemble method | Bag,<br>Gentle Boost,<br>Logit Boost,<br>AdaBoost,<br>RUS Boost | AdaBoost |
| Learner type | Decision tree | Decision tree |
| Number of Learners | 10-500 | 56 |
| Learning Rate | 0.001-1 | 0.92518 |
| Maximum number of splits | 1-974 | 12 |
| Number of predictors to sample | 1-15 | 15 |

**Table 4.** Optimized hyperparameters of the ensemble classification algorithm for breast cancer cell typing.

| Model Hyperparameters | range | Optimized Hyperparameters |
|---|---|---|
| Ensemble method | Bag,<br>AdaBoost,<br>RUS Boost | AdaBoost |
| Learner type | Decision tree | Decision tree |
| Number of Learners | 10-500 | 414 |
| Learning Rate | 0.001-1 | 0.99857 |
| Maximum number of splits | 1-3431 | 132 |
| Number of predictors to sample | 1-15 | 15 |

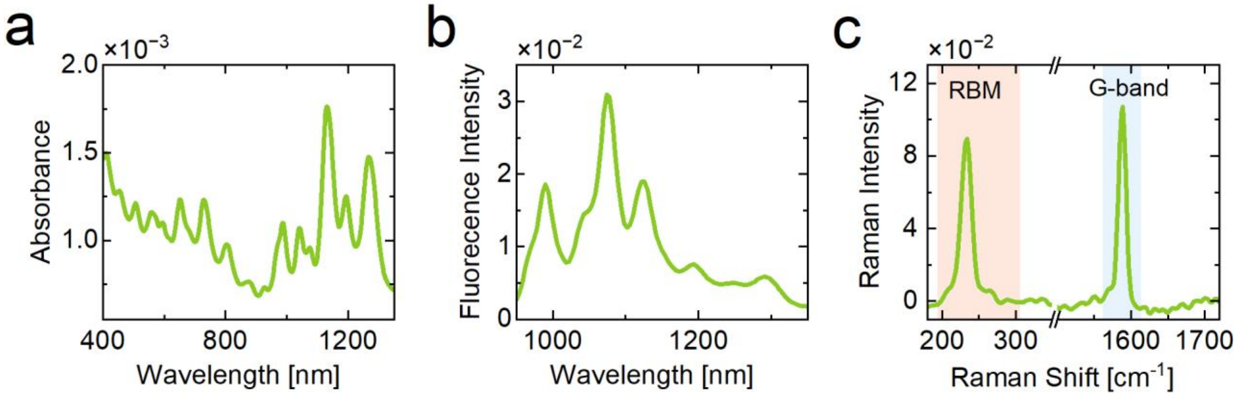


**Figure S1. Optical characterization of DNA-SWCNTs after functionalization.**
**(a)** UV–vis–NIR absorbance spectrum of DNA-functionalized SWCNTs, showing characteristic nanotube absorption features after dispersion. **(b)** Near-infrared fluorescence emission spectrum of DNA-SWCNTs, showing distinct emission peaks from multiple semiconducting SWCNT chiralities. **(c)** Raman Spectrum of DNA-SWCNTs, showing the characteristic peaks of the G band and RBM band.

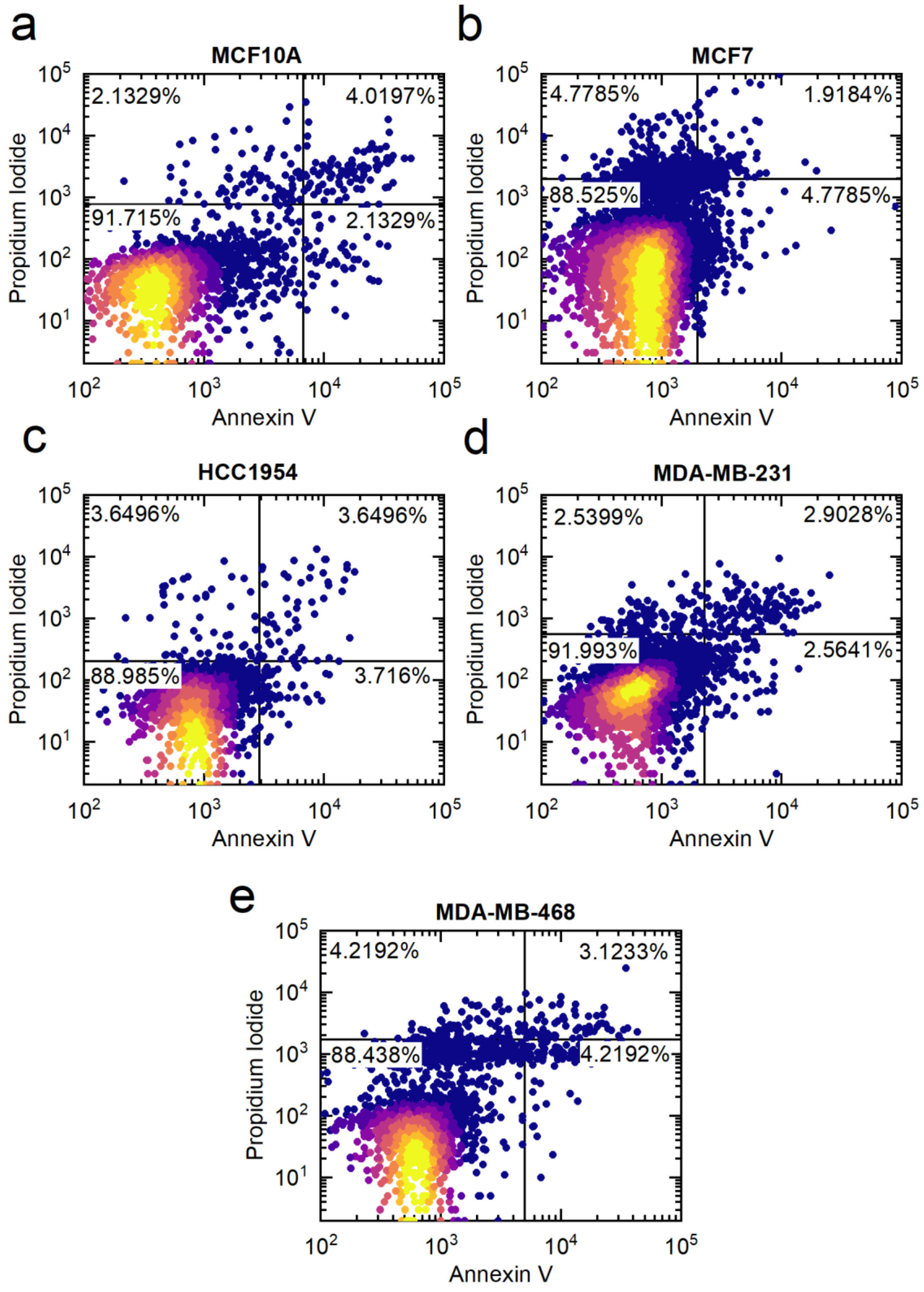


**Figure S2. Annexin V/propidium iodide viability analysis of untreated control cells.** Representative Annexin V and propidium iodide staining plots for untreated control cells from each breast epithelial or breast cancer cell line: **(a)** MCF10A, **(b)** MCF7, **(c)** HCC1954, **(d)** MDA-MB-231, and **(e)** MDA-MB-468. Quadrants indicate viable cells, early apoptotic cells, late apoptotic or necrotic cells, and dead cells based on Annexin V and propidium iodide fluorescence. These control measurements were used to establish cell-line-specific fluorescence gating thresholds for subsequent comparison with DNA-SWCNT-exposed samples.

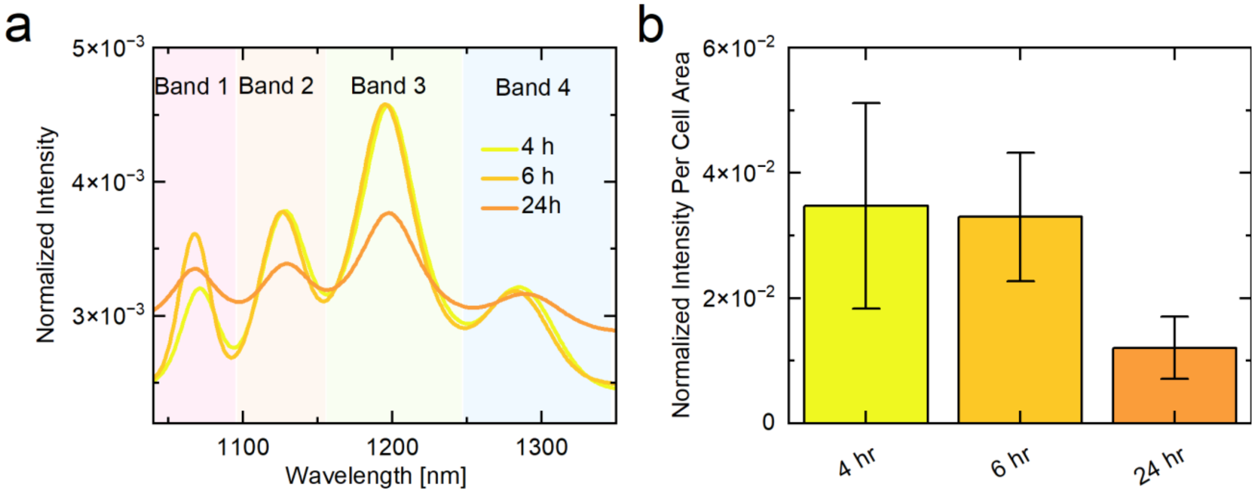


**Figure S3. Time-dependent DNA-SWCNT fluorescence response in MCF7 cells.**
**(a)** Normalized near-infrared fluorescence spectra from DNA-SWCNTs internalized by MCF7 cells after 4, 6, and 24 h nanotube exposure. Shaded regions indicate the spectral bands. **(b)** Area-normalized fluorescence intensity of MCF7 cells at 4, 6, and 24 h, showing reduced intracellular DNA-SWCNT fluorescence after 24 h compared with earlier time points. Error bars represent the variation across measured cells.

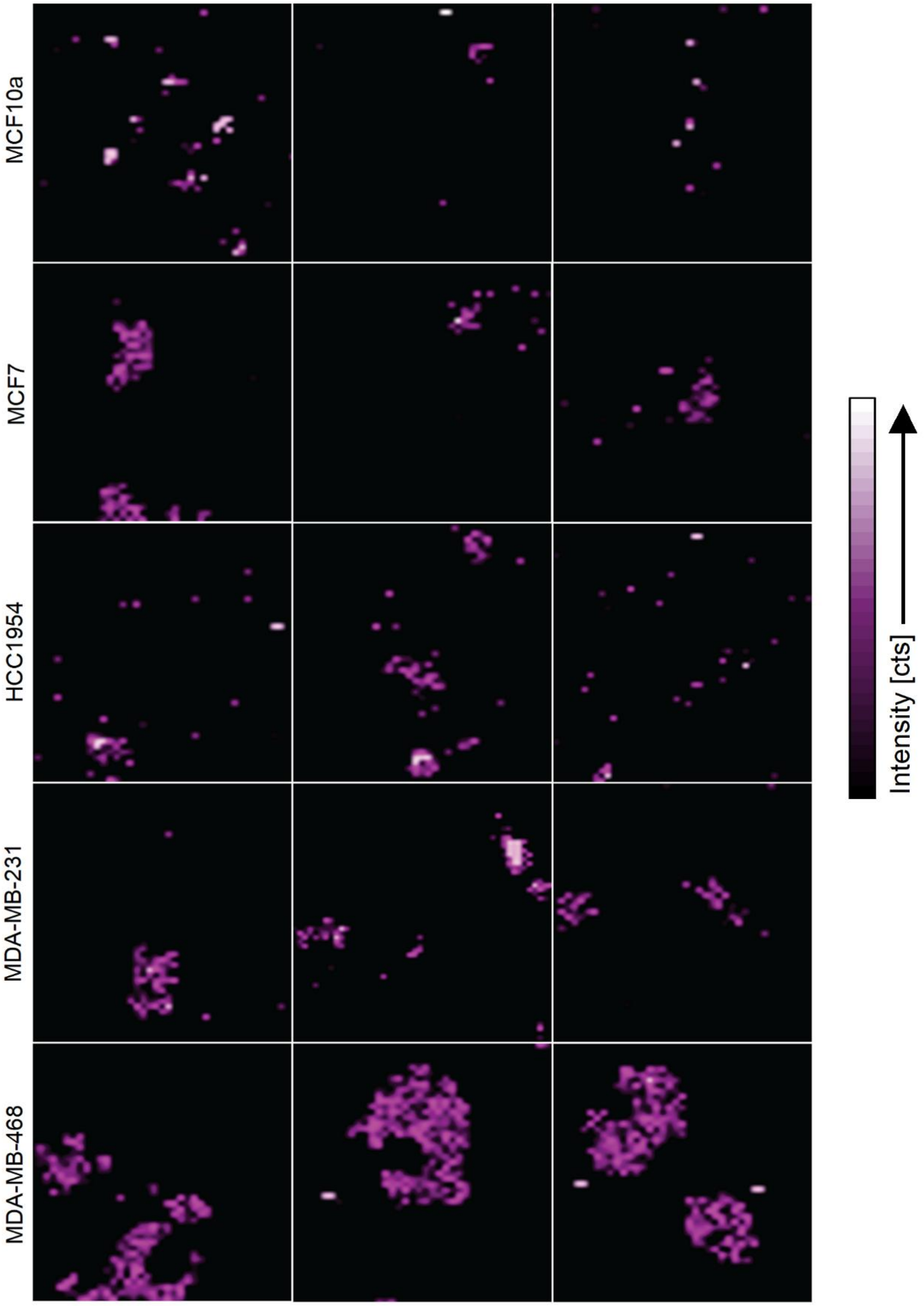


**Figure S4. Raman G-band intensity maps of DNA-SWCNT uptake across breast cell lines.**
Representative integrated Raman G-band intensity maps of DNA-SWCNTs in MCF10A, MCF7, HCC1954, MDA-MB-231, and MDA-MB-468 cells. Each row shows representative single-cell or cell-region maps for the corresponding cell line.

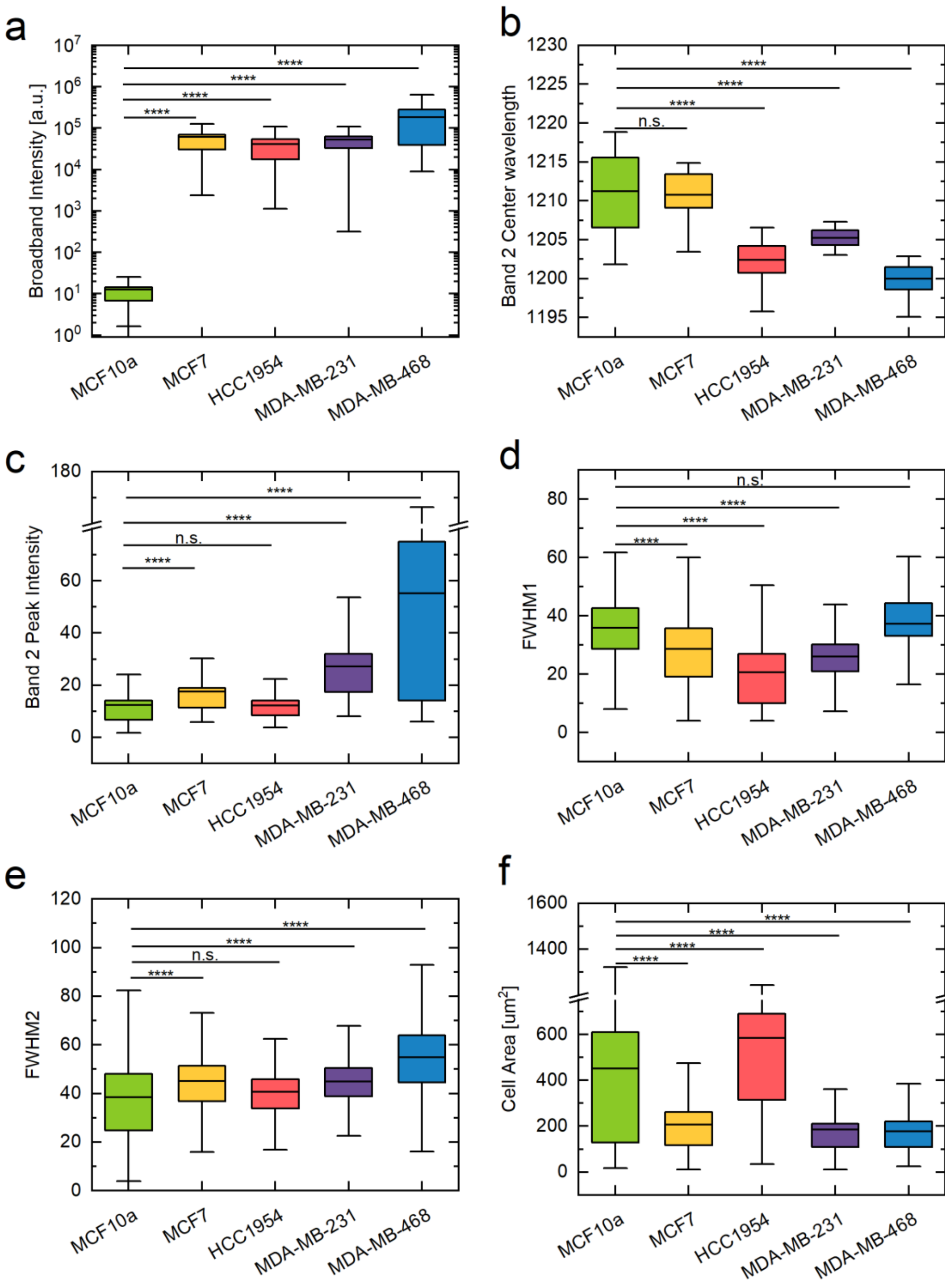


**Figure S5. Single-cell spectral and morphological features of different cell lines after DNA-SWCNT exposure.** Box plots of selected single-cell features extracted from DNA-SWCNT near-infrared fluorescence Spectra for MCF10A, MCF7, HCC1954, MDA-MB-231, and MDA-MB-468 cells: **(a)** broadband fluorescence intensity, **(b)** Band 2 center wavelength, **(c)** Band 2 peak intensity, **(d)** FWHM of Band 1, **(e)** FWHM of Band 2, and **(f)** cell area (n=200). Boxes indicate the 25th–75th percentile range, horizontal lines indicate the mean, and whiskers extend to the minimum and maximum values. Statistical significance was assessed using Kruskal–Wallis ANOVA followed by Dunn's multiple comparisons test. (****p < 0.0001 and n.s. indicates not significant)

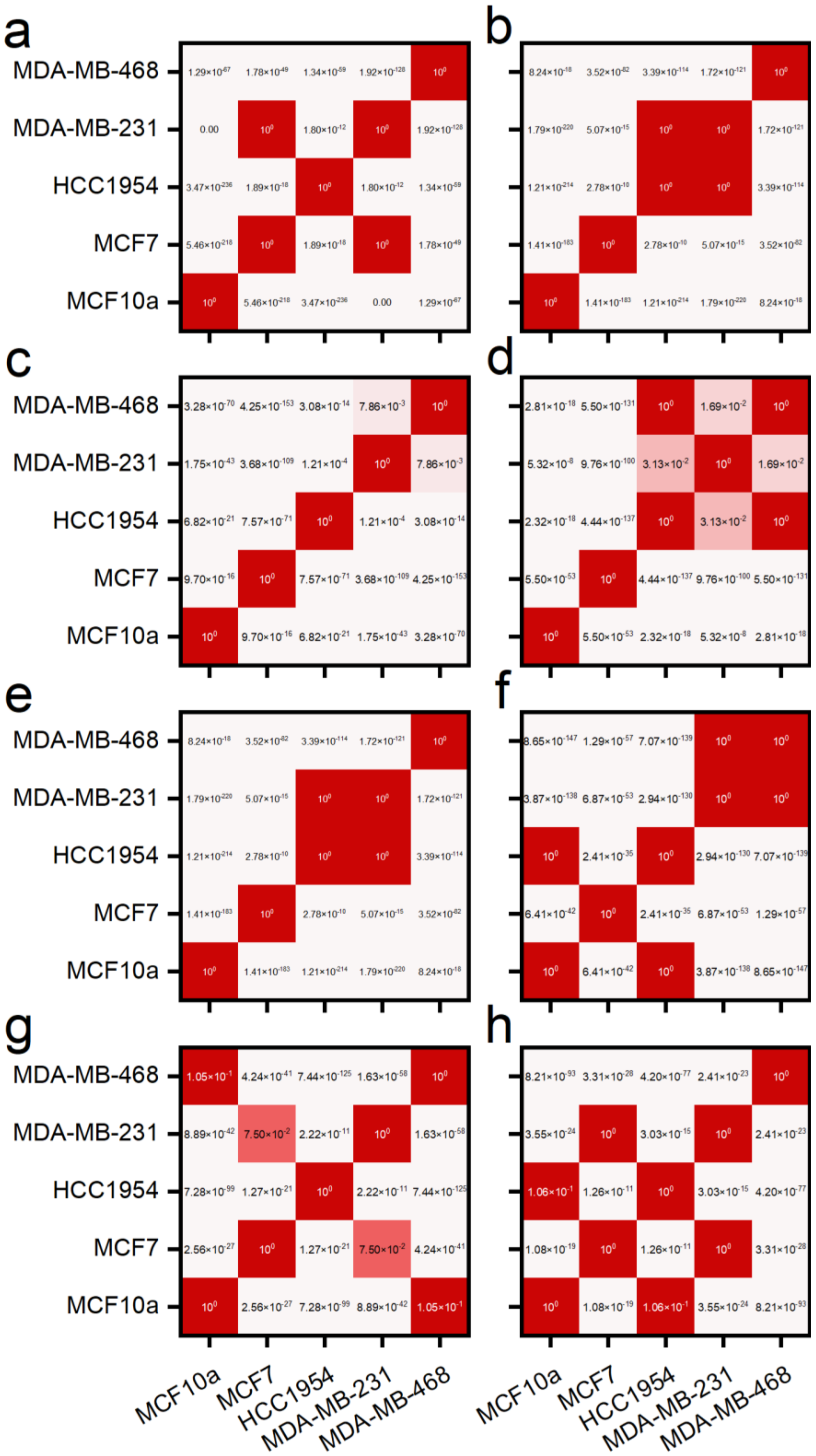


**Figure S6.** Pairwise statistical comparison of DNA-SWCNT spectral features across breast cell lines.
Pairwise comparison matrices showing adjusted p-values for selected single-cell DNA-SWCNT spectral features across MCF10A, MCF7, HCC1954, MDA-MB-231, and MDA-MB-468 cells. Features include (a) broadband intensity normalized by cell area, (b) Band 1/Band 2 intensity ratio, (c) Band 1 center wavelength, (d) Band 2 center wavelength, (e) Band 1 intensity, (f) Band 2 intensity, (g) FWHM of Band 1, and (h) FWHM of Band 2.

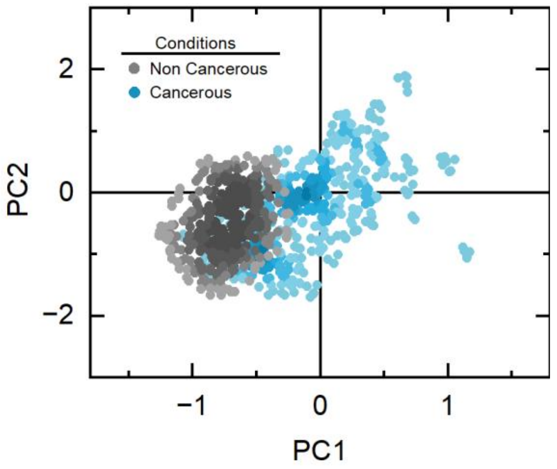


**Figure S7.** Principal component analysis of DNA-SWCNT spectral fingerprints from non-cancerous and cancerous breast cell models. Each point represents an individual cell projected onto the first two principal components.

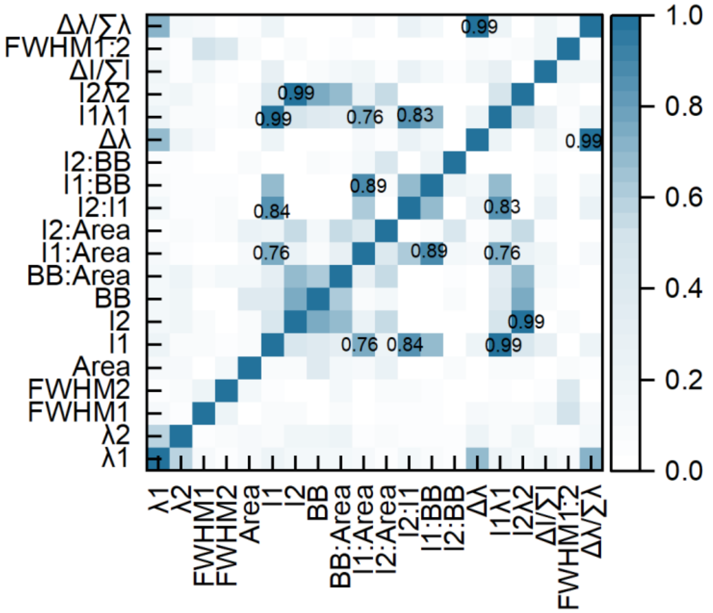

**Figure S8.** Pearson Correlation matrix of 20 DNA-SWCNT spectral features used for single-cell classification.

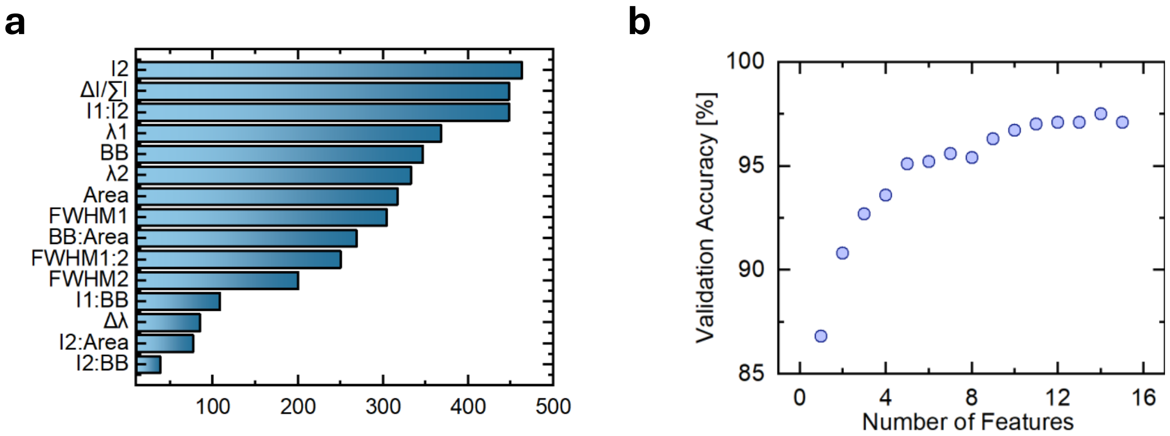


**Figure S9.** Feature ranking and classification performance as a function of the number of DNA-SWCNT spectral features. **(a)** Ranking of extracted spectral features based on Kruskal–Wallis analysis. **(b)** Classification accuracy of the breast cancer cell typing model as a function of the number of input sensor features. Ranked from most to least important.

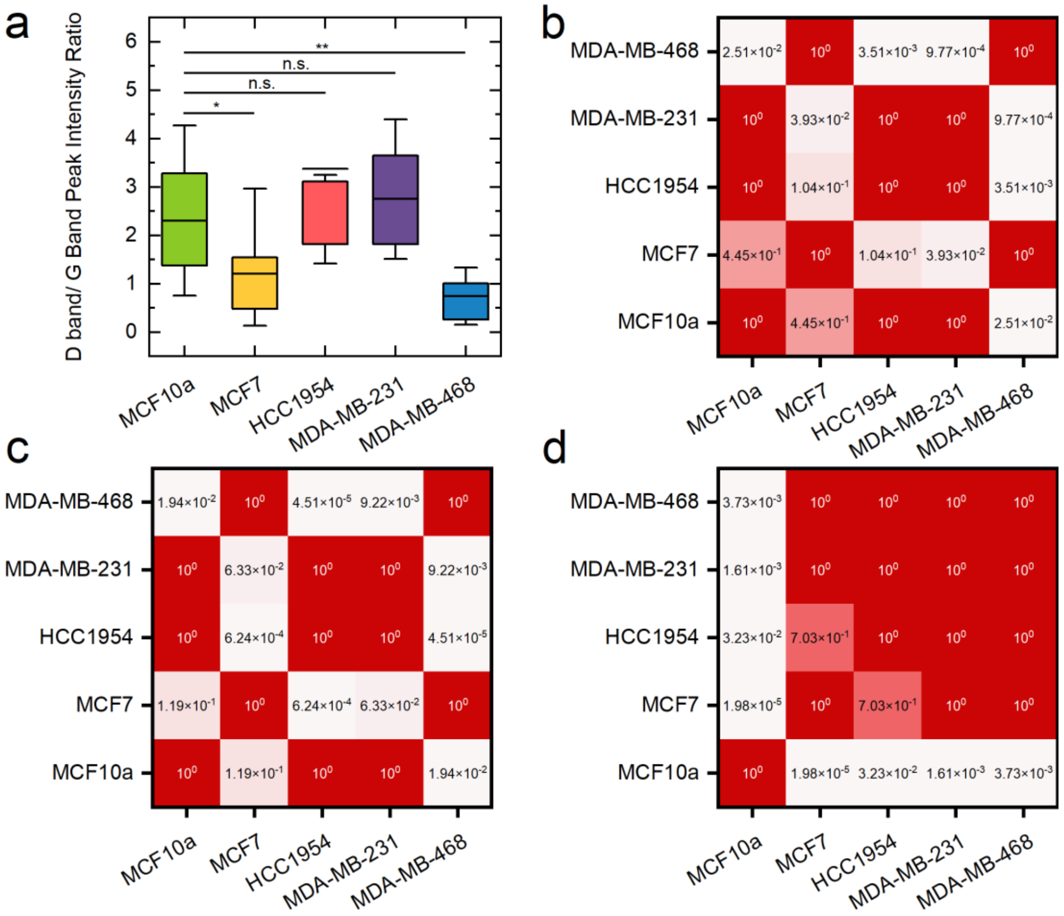


**Figure S10.** Confocal Raman microscopy analysis of SWCNT uptake by different cell types.
(a) Box plot showing the D-band/G-band peak intensity ratio measured from Raman spectra of DNA-SWCNT-exposed MCF10A, MCF7, HCC1954, MDA-MB-231, and MDA-MB-468 cells. Boxes indicate the 25th–75th percentile range, horizontal lines indicate the mean, and whiskers extend to the minimum and maximum values. Statistical significance was assessed using a Kruskal–Wallis ANOVA followed by Dunn's multiple comparisons test (*$p<0.05$ and **$p<0.01$). (b–d) Pairwise statistical comparison matrices for Raman-derived features across cell lines: (b) D/G band intensity ratio, (c)

integrated G-band intensity, and (d) RBM Band 2/Band 1 intensity ratio. Matrix values represent adjusted p-values for pairwise comparisons between cell lines.

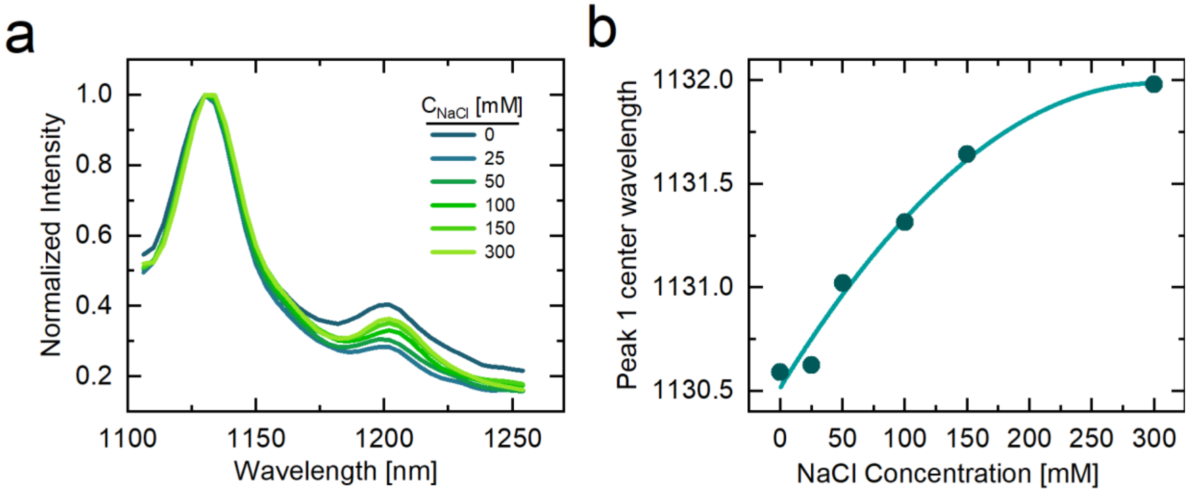


**Figure S11.** Solution-phase recapitulation of DNA-SWCNT spectral responses to ionic strength.
**(a)** Normalized near-infrared fluorescence spectra of DNA-SWCNTs in solution at increasing NaCl concentrations from 0 to 300 mM. **(b)** Peak 1 center wavelength as a function of NaCl concentration, showing a concentration-dependent red shift with increasing ionic strength.

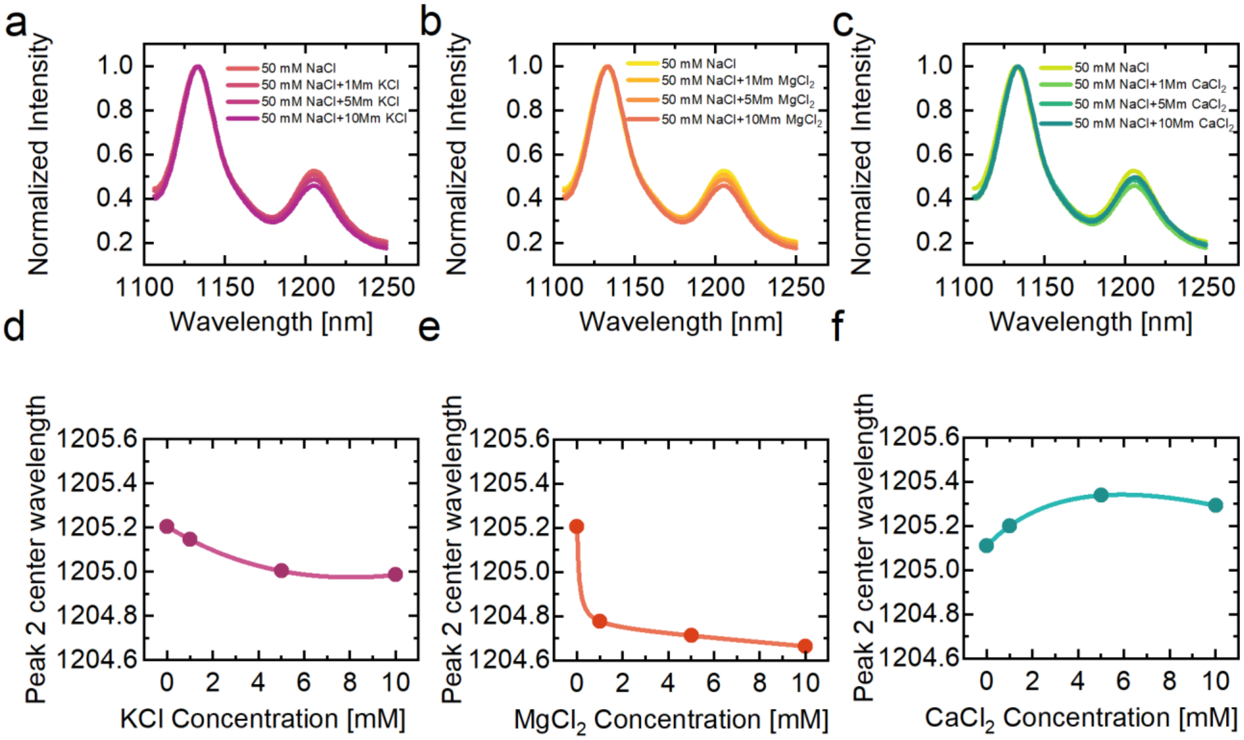


**Figure S12.** Solution-phase effects of intracellular ions on DNA-SWCNT fluorescence spectra.
**(a–c)** Normalized near-infrared fluorescence spectra of DNA-SWCNTs in 50 mM NaCl with increasing concentrations of **(a)** KCl, **(b)** $MgCl_2$, and **(c)** $CaCl_2$. **(d–f)** Corresponding Peak 2 center wavelength responses as a function of **(d)** KCl, **(e)** $MgCl_2$, and **(f)** $CaCl_2$ concentration. Monovalent and divalent ions produce distinct spectral responses, indicating that ionic composition can modulate DNA-SWCNT fluorescence and may contribute to cell-dependent spectral fingerprints observed in intracellular measurements.
